# Active fatty acid oxidation defines the cellular response towards reactive oxygen species

**DOI:** 10.1101/2020.07.13.200022

**Authors:** Lars Kaiser, Isabel Quint, René Csuk, Manfred Jung, Hans-Peter Deigner

## Abstract

Endocrine disrupting compounds (EDC) are ubiquitous in the human environment, displaying a highly relevant research topic. The impact of EDC on the differentiation of primitive cells, e.g. in hematopoiesis, is of particular interest. We found profound inhibitory effects of di-2-ethylhexyl phthalate (DEHP) on erythropoiesis and dendropoiesis, mediated via reactive oxygen species (ROS) generation. Neutrophil differentiation, however, was not affected by DEHP. ROS leads to a shift from glycolysis to the pentose phosphate pathway and diminishes ATP generation from glycolysis, ultimately resulting in apoptosis in both cell types. In neutrophils, ATP generation is held constant by active fatty acid oxidation (FAO), rendering these cells highly resistant against ROS. This relationship also holds true in HUVEC and HepG2 cells, also in combination with other organic peroxides. We, therefore, uncover a key mechanism for ROS quenching which further explains the distinct ROS quenching ability of different tissues.

## Introduction

Hematopoiesis is a fundamental process in the bone marrow of the human body, giving rise to all blood cells during each individuals lifetime (Doulatov et al., 2012). As there is a constant turnover, continuously replenishing the total blood cell pool, the hematopoietic system is uniquely vulnerable to changes in the environment (Laiosa and Tate, 2015). Importantly, endocrine disrupting compounds (EDC) are presumed to own a significant impact on hematopoiesis (Laiosa and Tate, 2015). EDC are defined as “exogenous chemicals or mixture of chemicals that interfere with any aspect of hormone action” and consist of a huge heterogenous group of compounds (e.g. dioxins, pesticides, phthalates, bisphenol A, etc.), found ubiquitously in the human environment (Lauretta et al., 2019; Zoeller et al., 2012). Therefore, assessment of effects of EDC on hematopoietic stem and progenitor cell (HSPC) differentiation is a highly relevant research topic in today’s society.

One compound from the class of EDC is di-2-ethylhexyl phthalate (DEHP), a plasticizer in polymer products, consisting of a benzene-dicarboxylic acid ring with two eight-carbon esters linked to it (Jo et al., 2011; Rowdhwal and Chen, 2018; Schaedlich et al., 2018). It is the most frequent used phthalate, especially in PVC-based medical devices (e.g. intravenous bags or infusion tubing), making up to 40% by weight of such materials (Erythropel et al., 2014). As DEHP is not covalently bound to the polymer matrix, leaching occurs into liquids in contact with the material, raising concerns about its impact on human health. As example, 50 to 70 mg/l DEHP have been found in blood bags, indicating high exposure levels for patients, especially those in intensive care units (Erythropel et al., 2014). Importantly, neonates at intensive care units have been identified as highly exposed, reaching DEHP levels up to 123.1 µg/ml in serum after transfusion exchange or 34.9 µg/ml after extracorporeal membrane oxygenation (Ernst et al., 2014). Therefore, most frequently tested DEHP concentrations under experimental conditions range between 0.1 to 100 µg/ml (Gutiérrez-García et al., 2019; Lee et al., 2018; Schaedlich et al., 2018; You et al., 2014). It should also further be noted, that DEHP is degraded into several metabolites, including mono-2-ethylhexyl phthalate (MEHP), which were shown to be even more toxic than DEHP itself.

Regarding the toxicity of DEHP, various effects have been observed in different model systems. Indeed, there are several reports on DEHP mediated effects in mice or rats, however, the sensitivity to DEHP-mediated toxicity, seems to be variable in different species (Jo et al., 2011; Shen et al., 2017; You et al., 2014; Zarean et al., 2016; Zhang et al., 2012). Furthermore, toxicity apparently differs between different cell types used. For example, DEHP was shown to impair fatty acid storage in human SGBS-adipocytes, decreases estradiol synthesis in human granulosa cells and reduces phagocytosis and secretion of several cytokines in differentiated THP-1 cells (Couleau et al., 2015; Ernst et al., 2014; Schaedlich et al., 2018). General toxicity, however, was absent in all cases. On the contrary, cytotoxic effects of DEHP were observed in rat ovarian granulosa cells, rat insulinoma cells, human hepatic stellate cells, mouse oocytes and human colon carcinoma cells (Amara et al., 2020; Gaitantzi et al., 2018; Lu et al., 2019; She et al., 2017; Tripathi et al., 2019). On the mechanistic level, several of the effects found seemed to be mediated via DEHP-dependent activation of the peroxisome proliferator-activated receptor gamma (PPARγ), or generation of reactive oxygen species (ROS) (Engel et al., 2017; Fang et al., 2019; Schaedlich et al., 2018; She et al., 2017; Tripathi et al., 2019; You et al., 2014). On the epidemiological level, phthalate exposure (including DEHP) has been linked with the occurrence of asthma and other allergic symptoms in children, also indicating some effect on human immune cells *in vivo* (Bornehag and Nanberg, 2010). In mice, this effect seems to be mediated via oxidative stress; nevertheless, modulation of dendritic cell differentiation derived from peripheral blood mononuclear cells (PBMC) by DEHP, was also observed (Ito et al., 2012; You et al., 2014). Therefore, an impact of DEHP on human hematopoiesis is a reasonable assumption. Indeed, DEHP was recently shown to reduce the *in vitro* expansion of HSC’s, also reducing the total number of derived total progenitors (Gutiérrez-García et al., 2019). Moreover, also a short time exposure (24 and 72h) of HSPC to DEHP was previously shown to selectively reduce colony formation (Manz et al., 2015). Given these facts, an impact of DEHP exposure on HSPC differentiation is evident; the exact mode of action of DEHP in modulating HSPC differentiation, however, is still not resolved.

Indeed, endocrine disrupting properties of DEHP may be responsible for the observed effects, as PPARγ activation has been shown to modulate hematopoiesis (Chute et al., 2010). Nevertheless, it also became clear during the recent years, that ROS are not merely harmful metabolic by-products, but are involved into various signalling pathways, which are strictly regulated (Bardaweel et al., 2018). In the context of hematopoiesis, ROS are known to be crucial for the “oxidative burst” in phagocytes, promote emergency granulopoiesis and define distinctive dendritic cell subset development, indicating their essential regulatory role during hematopoiesis (Kwak et al., 2015; Sheng et al., 2010; Yang et al., 2013). Likewise, quenching of ROS leads to inhibited erythroid differentiation from erythroid progenitors, further underlining the essential role of ROS during formation of distinct hematopoietic lineages (Nagata et al., 2007). Therefore, the observed effects of DEHP on hematopoiesis may also be mediated via ROS generation. Reduction of ROS is mainly driven by NADPH-dependent mechanisms, as NADPH is involved in glutathione and thioredoxin reduction (Snezhkina et al., 2020). Indeed, the main source for intracellular NADPH is the pentose phosphate pathway (PPP). However, isocitrate dehydrogenases (IDH), malic enzymes (ME) and NAD(P) transhydrogenases are recognized to be involved in NADPH regeneration as well (Aon et al., 2010; Fernandez-Marcos and Nóbrega-Pereira, 2016; Snezhkina et al., 2020). Of note, it is quite well known that cells can redirect glycolytic fluxes towards PPP, in order to reduce ROS levels via different regulatory mechanisms (Snezhkina et al., 2020). Glycolysis, however, is known to be the major energy source in various tissues, thus it is quite reasonable to expect that a prolonged redirection of fluxes towards PPP may possess negative effects on such tissues (Dagher et al., 2001).

We recently established and characterised a myeloid HSPC differentiation model, capable of differentiation into erythrocytes, dendritic cells and neutrophils. Between these lineages, metabolic fluxes uniquely differ, while neutrophils were the only lineage with active FAO (Kaiser et al., 2020). Here, we assessed the impact of DEHP on the different lineage differentiations and found profound inhibitory effects on erythroid and dendritic cell differentiation, whereas neutrophil differentiation was enhanced. Metabolic profiling indicated global disturbance of the lipidome, as well as reduced glutaminolysis and glycolysis in both negatively affected lineages. Mechanistic investigation revealed that effects of DEHP were mediated via enhanced ROS levels, rather than endocrine disruption of peroxisome proliferator-activated receptors (PPARs). Enhanced ROS levels, in turn, lead to diminished intracellular ATP concentrations, while NADPH levels were raised. As this relation was absent in neutrophils, we investigated the relation between FAO activity and ROS quenching ability and found that active FAO is essential to combat prolonged oxidative stress. As this relationship could also be verified in two additional cell lines differing in their activity of FAO, we propose ROS quenching by FAO activity to be a general mechanism.

## Results

### DEHP induces apoptosis in erythrocytes and dendritic cells, while enhancing neutrophil maturation

As large quantities of all blood cells are formed from just few progenitor cells, large expansion of cell numbers accompanies hematopoietic lineage differentiation. We therefore initially assessed the proliferation rates of the different lineages in presence of 25.6 µM (10 µg/ml), 128.2 µM (50 µg/ml), 256.41 µM (100 µg/ml) and 641 µM (250 µg/ml) DEHP. Raising DEHP concentrations significantly lowered the proliferation rates during erythroid and dendritic cell differentiation in a concentration-dependent manner (Fig. 1 a, i). Proliferation rates during neutrophil lineage formation, however, remained unaffected (supplemental Fig. S1 a). Furthermore, surface markers of early (CD71) and late (CD235a) erythroid differentiation, as well as *HBB* expression, were drastically reduced by DEHP at day six in erythrocytes (Fig. 1 c-h) (Marsee et al., 2010). The substantially lowered HBB expression is also clearly supported by visual control of cell pellets from the resulting cell populations, as DEHP treated populations largely lack reddish colour, resulting from iron haemoglobin complexes (Fig. 1 b). In contrast, *GPNMB* expression, which is related to antigen presenting cells (APC), was increased in dendritic cells by DEHP (Fig. 1 j) (Gutknecht et al., 2015). As *GPNMB* expression is also found in monocytes/macrophages, we assessed the expression of CD14, which was unaffected by treatment with 256.41 µM DEHP (Fig. 1 k,l). Altered *GPNMB* expression, therefore, seems not to be related to a shift towards monocytic differentiation (Pahl et al., 2010). Indeed, reduced proliferation in erythrocytes and dendritic cells apparently result from increased apoptosis, as caspase 3/7 activity was increased by DEHP (Fig. 1 m,n). In line with the proliferation rates, no change of caspase 3/7 activity in the presence of DEHP was observed in neutrophils (results not shown). Expression of functional neutrophil lineage related genes *ELANE* and *S100A8*, however, were reduced by the addition of DEHP (Fig. 1 p, q). Expression of both genes in fact was shown to be maturation stage dependent; *ELANE* expression decreases at the stage of metamyelocytes, while expression of *S100A8* increases in metamyelocytes and decreases again in mature polymorphonuclear neutrophils (Grassi et al., 2018). This expression pattern, therefore, indicates either severe inhibition of neutrophil differentiation or increased maturation by DEHP. As the proportion of CD14^low^ cells, representing more mature neutrophils, is increased by DEHP, the latter assumption seems to hold true (Fig. 1 o) (Antal-Szalmas et al., 1997; Grassi et al., 2018).

**Figure 1.**
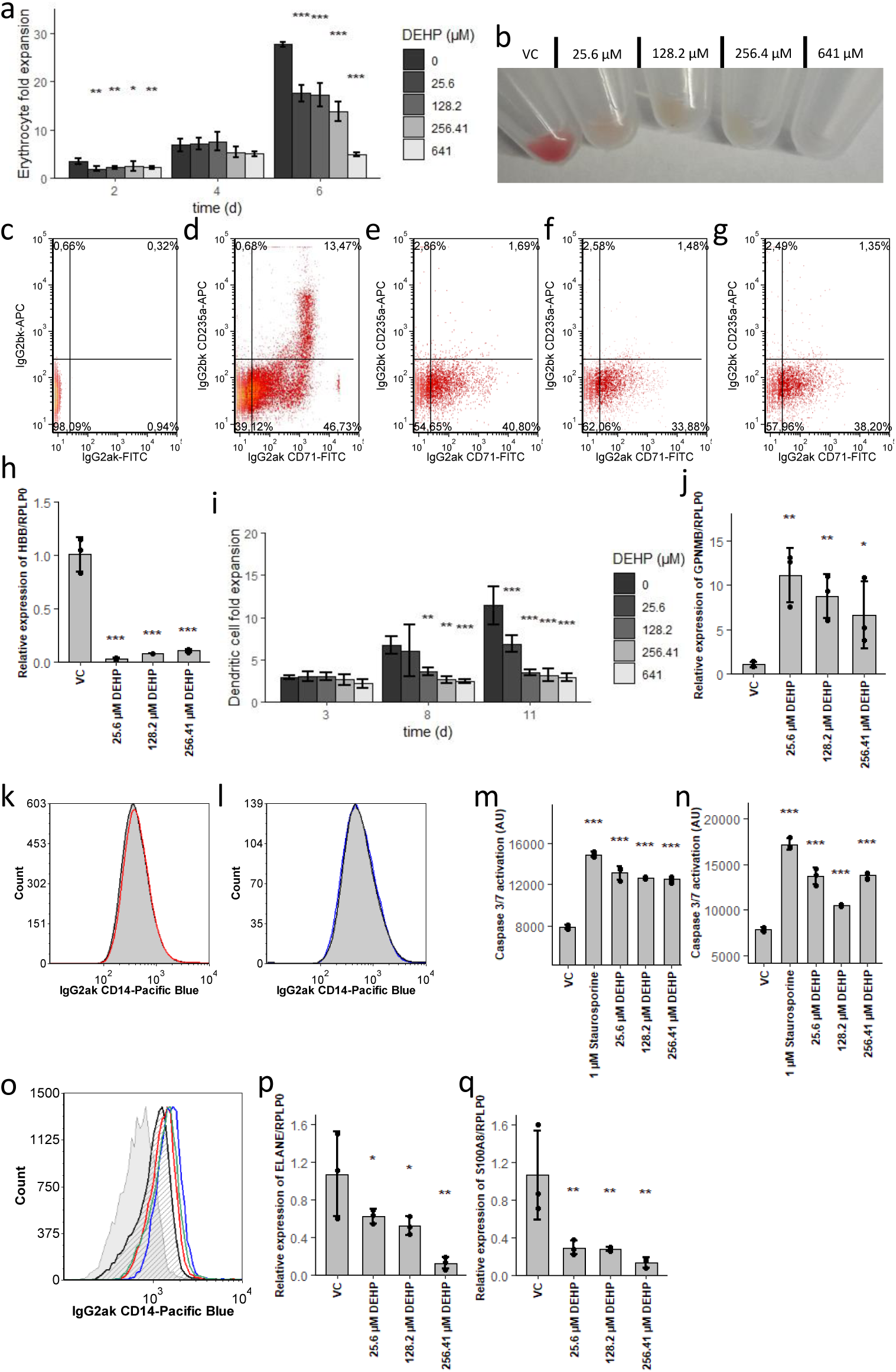
Differential disturbance of myeloid lineage differentiation by DEHP. See also Figure S1. (a,i) Expansion rate of erythrocytes (a) and dendritic cells (i), treated with raising concentrations of DEHP. (b) Cell pellet of erythroid populations after 6 days, treated with different concentrations of DEHP. (c-g) CD71 and CD235a expression of erythroid populations after 6 days, treated with different concentrations of DEHP. From left to right: isotype control (c), vehicle control (d), 25.6 (e), 128.2 (f) and 256.41 µM DEHP (g). (h) Impact of rising concentrations of DEHP on HBB/RPLP0 expression of erythrocytes after 6 days. (j) Impact of DEHP on GPNMB/RPLP0 expression of dendritic cells. (k,l) Presence of CD14 on dendritic cells, treated with 256.41 µM DEHP (l), compared to vehicle control (k). (m,n) Caspase activity in erythroid populations after 2 days (m) and in dendritic cell populations after 3 days (n), treated with different concentrations of DEHP. Staurosporine was used as positive control. (o) CD14 expression of neutrophils treated with 25.6 µM (red), 128.2 µM (green) and 256.41 µM DEHP (blue), compared to vehicle control (black) and isotype control (grey). (p,q) Impact of DEHP on ELANE/RPLP0 (q) and S100A8/RPLP0 expression (r) in neutrophils. Data are represented as mean ± SEM. Significance was assessed by one-way ANOVA (*p<0.05; **p<0.01; ***p<0.001).

Taken together, our data demonstrates that DEHP selectively disturbs erythropoiesis and dendropoiesis via induction of apoptosis. Neutrophil granulopoiesis, however, seems to be enhanced by DEHP.

### DEHP alters lipidome composition and lowers glycolysis, glutaminolysis and polyamine synthesis in erythrocytes

Since DEHP displayed differential effects on myeloid differentiations, we sought on identifying the mechanisms behind these opposing actions. As the metabolism plays a central role during hematopoietic lineage commitment and the cellular response towards exogenous stimuli, we performed quantitation of 200 metabolites over the differentiation period, including amino acids and biogenic amines, acylcarnitines, glycerophospho-(GPL) and sphingolipids and the sum of hexoses (Kaiser et al., 2020; Oburoglu et al., 2014, 2016).

As shown in Fig. 2 a-c, we found in the erythroid populations a total of 92 metabolites at day 2, 51 metabolites at day 4 and 55 metabolites at day 6 were significantly altered by the different DEHP concentrations. During the time course, the majority of significant acyl-acyl phosphatidylcholines (PC; “PC.aa.” species) and sphingolipids (SL; “SM” “Ceramide” and “Sph” species) were raised at day 2, while found to be lowered at day 4 and day 6 (supplementary table 1-3). Additionally, several ether phosphatidylcholines (EL; “PC.ae.” species) increased at day 2 and 6, while being lowered at day 4 (supplementary table 1-3). Furthermore, several lysophosphatidylcholines (lysoPC; “lyso.PC.a.” species) showed elevated concentrations at all time points. As we previously identified high fatty acid (FA) synthesis in combination with low FA oxidation (FAO) as underlying mechanism for increased glycerophospholipid and SL levels in erythrocytes, it is reasonable to assume initial stimulation of FA and GPL synthesis by DEHP (Kaiser et al., 2020). This is further supported by the sum of measured ester GPL and ether GPL, which were raised at day 2 by DEHP (Fig. 2 d,e). Raised levels of several lysoPC species during the time course, as well as lowered SL from day four on, indicate enhanced degradation of PC and SL in later stages. As phospholipases for ester glycerophospholipids (GPL) often can hardly hydrolyse corresponding ether GPL species, EL remain unaffected from this enhanced degradation (Uyama et al., 2012). Nevertheless, as acylcarnitines, reflecting cellular total acyl-CoA and thus free FA levels, were not altered by DEHP, reduced FA synthesis seems to be present as well.

**Figure 2.**
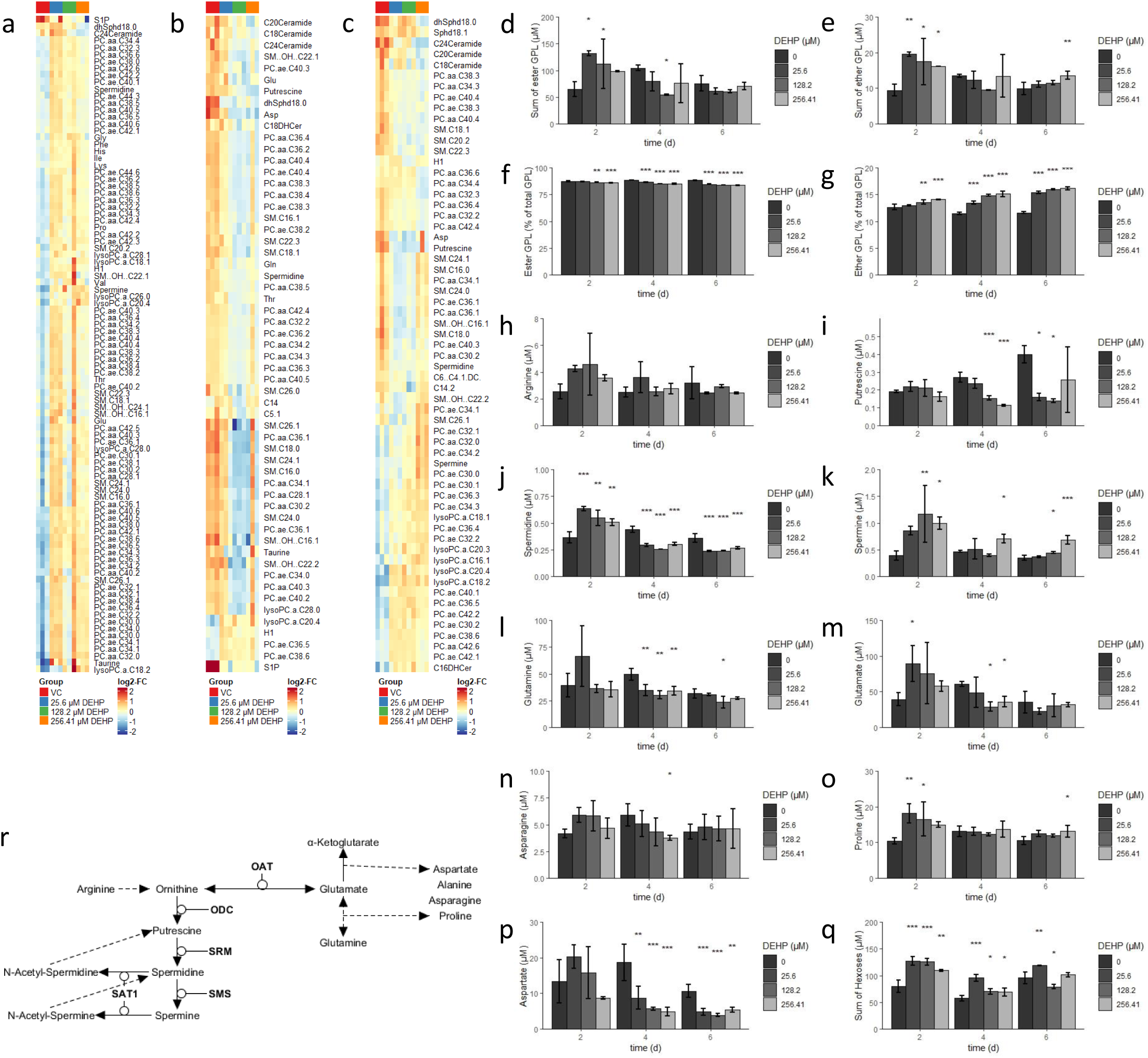
DEHP alters the lipidome and supresses glycolysis, glutaminolysis and polyamine synthesis in erythrocytes. (a-c) FDR-significant altered metabolites by raising DEHP concentrations during erythroid differentiation after 2 days (a), 4 days (b) and 6 days (c). See also Table S1-3. (d) Sum of ester glycerophospholipids (GPL) in erythroid populations, treated with raising concentrations of DEHP. (e) Sum of ether glycerophospholipids (GPL) in erythroid populations, treated with raising concentrations of DEHP. (f) Percentual content of ester glycerophospholipids in erythroid populations, treated with increasing concentrations of DEHP. (g) Percentual content of ether glycerophospholipids in erythroid populations, treated with raising concentrations of DEHP. (h-q) Concentration of arginine (h), putrescine (i), spermidine (j), spermine (k), glutamine (l), glutamate (m), asparagine (n), proline (o), aspartate (p) and sum of hexoses (q) altered by increasing DEHP concentrations during erythroid differentiation. (r) Model depicting the interconnection of altered metabolites, displayed in (h-p). Data are represented as mean ± SEM. Significance was assessed by one-way ANOVA (*p<0.05; **p<0.01; ***p<0.001).

We further found enhanced levels of hexoses at day 2 and 4, indicating that catabolism of glucose is lowered by DEHP. In addition, higher levels of glutamate, proline, spermidine and spermine at day 2 (Fig. 2 m,j,k,o) indicate a shift from glutamate conversion to α-ketoglutarate towards anabolism of other products, including polyamines (model Fig. 2 r). Lowered glutamine uptake later than day 4 (Fig. 2 l), in turn, seems to reduce levels of the related metabolites putrescine, spermidine, glutamate, asparagine and aspartate (Fig. 2 I,j,m,n,p). Taken together, DEHP seems to enhance GPL, EL and SL synthesis until day 2, while lowering GPL and SL levels afterwards and supresses glycolysis, glutaminolysis and polyamine synthesis.

### DEHP lowers glycerophospholipid and sphingolipid synthesis, glycolysis and glutaminolysis in dendritic cells

In the case of dendritic cell differentiation, we found a total of 5 metabolites altered by DEHP at day 3, while 115 and 99 were altered at day 8 and 11, respectively (Fig. 3 a-c, also supplementary table 4-6). During the time course, the majority of PC, EL and SL were lowered in a concentration dependent manner from day 8 on. This is also reflected by the sum of the measured ester GPL and ether GPL, which were both found remarkably lowered by DEHP (Fig. 3 d,e). The percentual content of ester and ether lipids, however, was not substantially altered by DEHP (Fig. 3 f,g). Taken together, the data indicate a lower synthesis of GPL and SL, as the majority of assessed lysoPCs were lowered by DEHP. Since DEHP seems to lower glycolysis, glutaminolysis and polyamine synthesis in erythrocytes, we also focussed on these metabolic pathways in dendritic cells. Indeed, the sum of hexoses was again significantly raised by DEHP, also indicating a lower rate of glycolysis in dendritic cells (Fig. 3 o). Furthermore, putrescine, spermidine, spermine, glutamate and aspartate levels were lowered by DEHP at day 8 and 11, indicating a lower rate of glutaminolysis and polyamine synthesis (Fig. 3 i-l,n). In contrast, arginine and proline levels were raised by DEHP, indicating a diminished conversion of arginine to ornithine, as well as enhanced conversion of glutamate to proline (Fig. 3 h,m). Overall, these results again indicate a lower rate of glycolysis, glutaminolysis and polyamine synthesis, caused by the increasing DEHP concentrations. Indeed, in dendritic cells, the total lipid anabolism seems to be inhibited, a fact which was not observed in erythrocytes. The total acyl-CoA pool controlling the rate of GPL, EL and SL synthesis, however, is fuelled by cholesterol ester (CE) and triglyceride (TG) degradation in dendritic cells, while *de novo* lipid synthesis replenishes acyl-CoA levels in erythrocytes (Kaiser et al., 2020). These different sources for lipid anabolism, in turn, may be responsible for the differential effects of DEHP on the lipidome of both lineages. DEHP, therefore, lowers total lipid levels, glycolysis, glutaminolysis as well as polyamine synthesis in dendritic cells.

**Figure 3.**
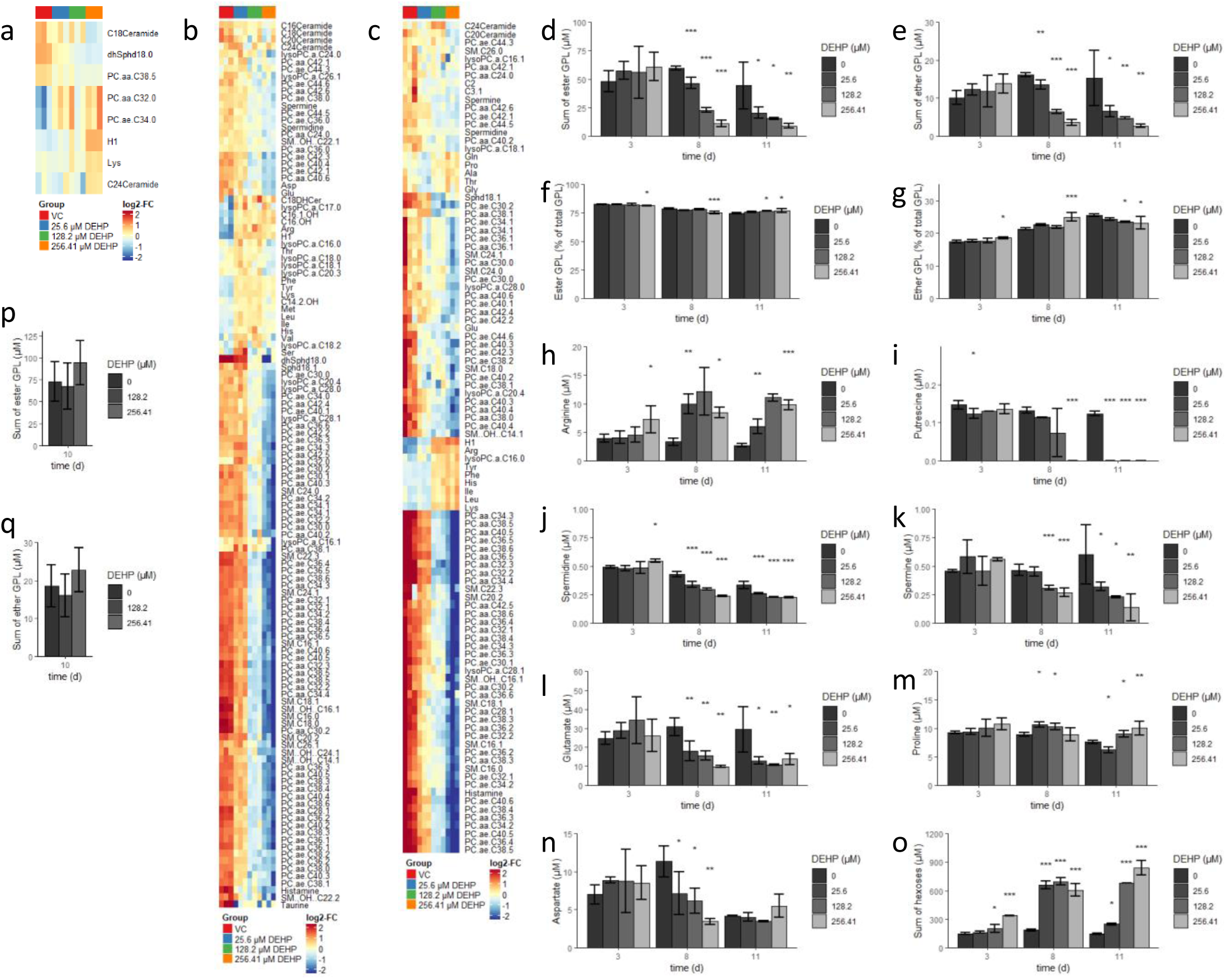
DEHP supresses glycerophospholipid and sphingolipid synthesis, glycolysis, glutaminolysis and polyamine synthesis in dendritic cells. (a-c) FDR-significantly altered metabolites in response to increasing DEHP concentrations during dendritic cell differentiation after 3 days (a), 8 days (b) and 11 days (c). See also Table S4-6. (d) Sum of measured ester glycerophospholipids (GPL) in dendritic cell populations, treated with rising concentrations of DEHP. (e) Sum of ether glycerophospholipids (GPL) dendritic cell populations, treated with increasing concentrations of DEHP. (f) Percentual content of ester glycerophospholipids in dendritic cell populations, treated with rising concentrations of DEHP. (g) Percentual content of ether glycerophospholipids in dendritic cell populations, treated with increasing concentrations of DEHP. (h-o) Concentrations of arginine (h), putrescine (i), spermidine (j), spermine (k), glutamate (l), proline (m), aspartate (n) and sum of hexoses (o) altered by distinct DEHP concentrations during dendritic cell differentiation. (m) Sum of ester glycerophospholipids (GPL) determined in neutrophil populations, treated with increasing concentrations of DEHP. (n) DEHP-concentration dependent sum of ether glycerophospholipids (GPL) in neutrophil populations, treated with raising concentrations of DEHP. Data are represented as mean ± SEM. Significance was assessed by one-way ANOVA (*p<0.05; **p<0.01; ***p<0.001).

In contrast, we did not find any significantly altered metabolites by 128.2 (50 µg/ml) and 256.41 µM (100 µg/ml) DEHP in neutrophils within 10 days, also the sum of measured ester GPL and ether GPL remained unchanged (Fig. 3 p,q). As major metabolic alterations by DEHP were observed in the lipidome of both other lineages and neutrophils are the only lineage with active FAO, we conclude a relationship between DEHP and FAO activity (Kaiser et al., 2020; Oburoglu et al., 2016; Riffelmacher et al., 2017).

### The effect of DEHP is not mediated via peroxisome proliferator-activated receptors or via activation of FAO

Lowering of GPL and SL levels can be caused by raised catabolic or lowered anabolic reactions. As increasing degradation of GPL and SL would result in higher levels of available free fatty acids, this would also be reflected by an increase in acylcarnitine concentrations (McCoin et al., 2015). However, the majority of acylcarnitines measured was not changed during the time course, implicating either enhanced degradation via FAO or reduced synthesis of corresponding CoA (and thus carnitine) species. As the regulation of FAO is implicated in HSPC fate decision, we tested the hypothesis that DEHP disturbs lipid metabolism via upregulation of FAO, leading to the observed selective toxic effects (Kaiser et al., 2020; Oburoglu et al., 2016).

Typically, the effects mediated by DEHP are attributed to PPARγ activation or activation of PPARα by its metabolite MEHP (Engel et al., 2017; Fang et al., 2019; Petit et al., 2018; Schaedlich et al., 2018). PPAR’s are a group of nuclear receptors, which regulate nutrient-dependent transcription (Nagasawa et al., 2005). Three isoforms of PPAR have been identified, namely α, β/d and γ, differently affecting lipid metabolism. While PPARα activation has been mainly associated with increased FAO, PPARγ activation has been associated with increased lipid storage and glucose homeostasis (Ahmed et al., 2007; Dubois et al., 2017; Lamichane et al., 2018; Tyagi et al., 2011; Wang, 2010). We, therefore, tested if the effect of DEHP is mediated via enhanced FAO, activation of PPARα or PPARγ, by using the antagonists etomoxir for CPT1a, GW-9662 for PPARγ and GW-6471 for PPARα. None of the antagonists used, however, were able to restore proliferation rates or gene expression patterns in both cell lines, even though corresponding positive controls confirmed the inhibitory effect of PPARα/γ activation in both lineages (supplemental Fig. S1 b-i). We therefore conclude that the observed lineage selective toxic effect of DEHP is neither mediated via activation of PPAR’s, nor by enhancing FAO.

### DEHP induces a ROS-mediated shift from Glycolysis to PPP, which is fatal for cell lineages without active FAO

As DEHP-induced lineage selective effects were apparently not induced via modulation of PPAR signalling or FAO activity, we focussed on the other prominent mode of action of DEHP, namely ROS generation. In fact, we found strongly increased levels of oxidative stress by DEHP treatment in erythrocytes and dendritic cells, while neutrophils displayed only a slight increase (Fig. 4 a-c). Apparently, this increase in ROS is also accompanied with a considerable decrease in ATP levels, in neutrophils, on the contrary, ATP levels were increased by DEHP (Fig. 4 d-f). Fuelling of the TCA and thus ATP production, was previously shown to be mediated via glycolysis and glutaminolysis in erythrocytes and dendritic cells (Kratchmarov et al., 2018; Oburoglu et al., 2014). In neutrophils, however, FAO is the main source for the TCA (Kaiser et al., 2020; Riffelmacher et al., 2017). As a shift from glycolysis to PPP in response to ROS is quite well-known, we presumed that lowered ATP levels are due to this effect (Snezhkina et al., 2020). In fact, we have found considerably raised concentrations of NADPH upon DEHP treatment in erythrocytes and dendritic cells (Fig. 4 g,h), supporting this interpretation. As this results in lower activity of glycolysis, concomitantly leading to lower substrate availability for TCA, this ultimately reduces ATP production. Lower ATP generation in turn, again lowers glucose phosphorylation by hexokinases, resulting in lower substrate availability for PPP, thus leading to a fatal loop, ultimately resulting in cell death. This assumption is also supported by the observed increase in hexoses, upon DEHP treatment (Fig. 2 q and Fig. 3 o). In addition, this mechanism further explains several other observed metabolic alterations. In erythrocytes for example, DEHP initially increases GPL and SM levels likely due to higher NADPH concentrations, as erythrocytes preferably perform lipid *de novo* synthesis and elongation, both dependent on NADPH (Kaiser et al., 2020). As a result of the fatal loop, NADPH levels are drastically lowered at later stages, resulting in disturbance of both anabolic processes, in consequence leading to lower levels of lipids containing long-chain fatty acids. On the contrary, GPL and SM levels in dendritic cells are drastically decreased at all time points due to DEHP treatment. This, however, again appears to be related to initial metabolic fluxes, as dendritic cells were shown to generate both from lipolysis of cholesterol esters and triglycerides, following CoA activation of free fatty acids, preferably by ACSL1 (Kaiser et al., 2020). As the latter is ATP dependent, the DEHP-mediated decrease in total ATP ultimately results in the observed decrease of GPL and SM due to lowered substrate availability (Soupene and Kuypers, 2008). In addition, several other metabolites, known to exhibit antioxidant activity (e.g. Taurine or polyamines), in erythrocytes and dendritic cells, are reduced at later stages, also indicating increased oxidative stress (Ripps and Shen, 2012; Snezhkina et al., 2020).

**Figure 4.**
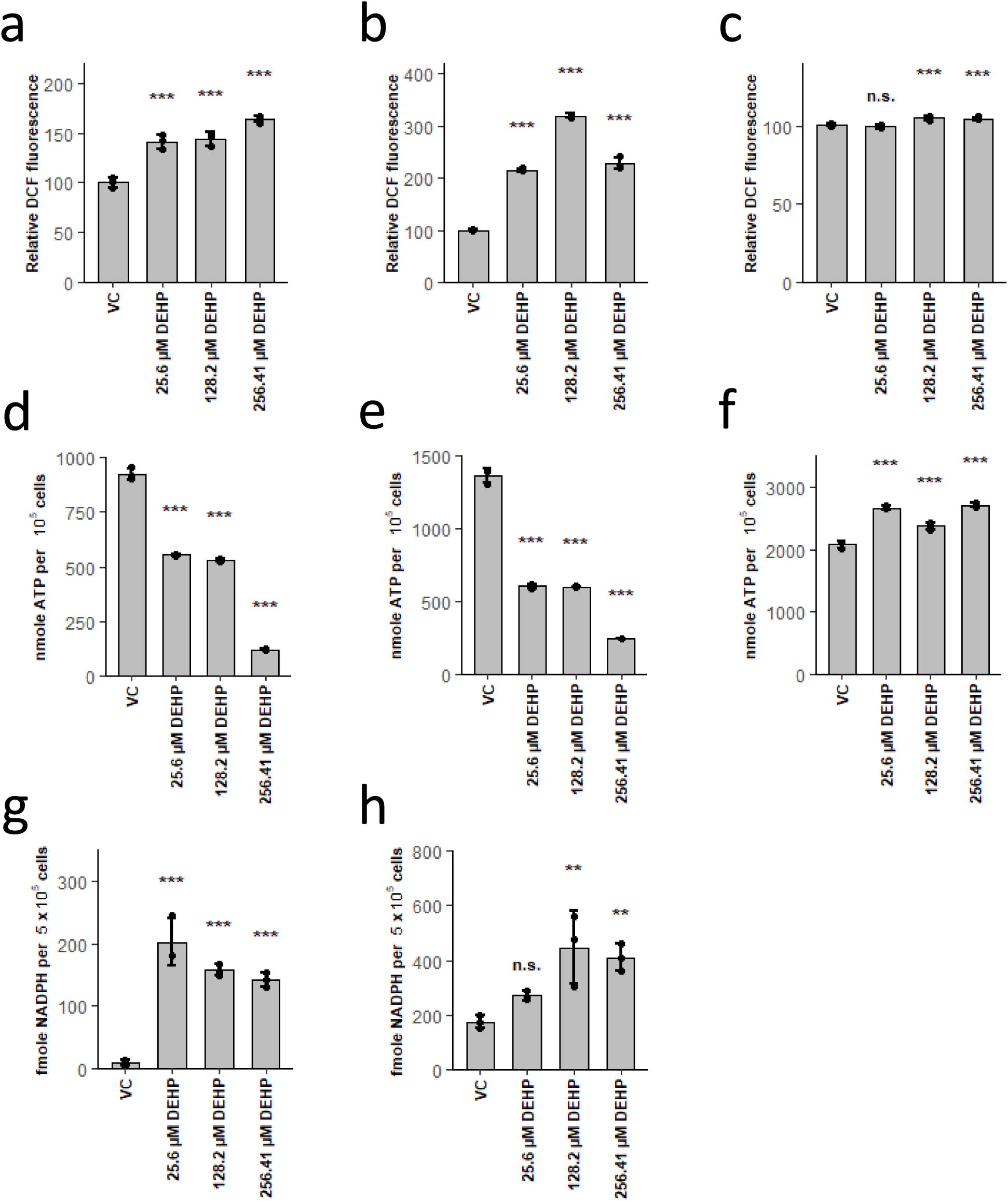
DEHP induces ROS, leading to a shift from glycolysis to PPP, resulting in ATP depletion in erythrocytes and dendritic cells. (a-c) Relative DCF fluorescence after treatment with DEHP in erythrocytes for 4h (a), dendritic cells for 3d (b) and neutrophils for 3d (c). (d-f) ATP levels in erythrocytes after 2d (d), dendritic cells after 3d (e) and neutrophils after 3d (f) treatment with DEHP. (g,h) NADPH levels in erythrocytes after 2d (g) and dendritic cells after 3d (h) treatment with DEHP. Data are represented as mean ± SEM. Significance was assessed by one-way ANOVA (*p<0.05; **p<0.01; ***p<0.001, n.s. not significant).

In neutrophils, however, no effects on the assessed metabolome were observed. Indeed, DEHP did merely induce a slight increase of oxidative stress, ATP levels were not lowered as well, on the contrary, increased. Even though increased flux via PPP due to increased oxidative stress should also occur in neutrophils, it seems like neutrophils can easily compensate ATP levels. As neutrophils were the only lineage with active FAO, the relation to lowered increase in ROS, in combination with stable ATP levels, seems obvious. As ATP is preferably generated via FAO, shifting fluxes from glycolysis to PPP here does not result in the fatal loop, observed in both other lineages. This interpretation is further supported by results, obtained in tumour cells; in these cells inhibition of FAO via etomoxir impairs NADPH production, increases ROS and decreases ATP, resulting in cell death (Pike et al., 2011).

In order to verify the essential role of FAO in ROS clearance, we pre-treated differentiating neutrophils with the CPT1a inhibitor etomoxir, following DEHP treatment. In line with our hypothesis, significantly higher oxidative stress was induced by DEHP if FAO is inhibited (Fig. 5 a). Furthermore, dichlorofluorescein (DCF) fluorescence remained low, when antioxidants N-acetylcysteine (NAC) or butylated hydroxyanisole (BHA) were added to etomoxir pre-treatment, verifying that increased fluorescence is due to higher oxidative stress levels.

**Figure 5.**
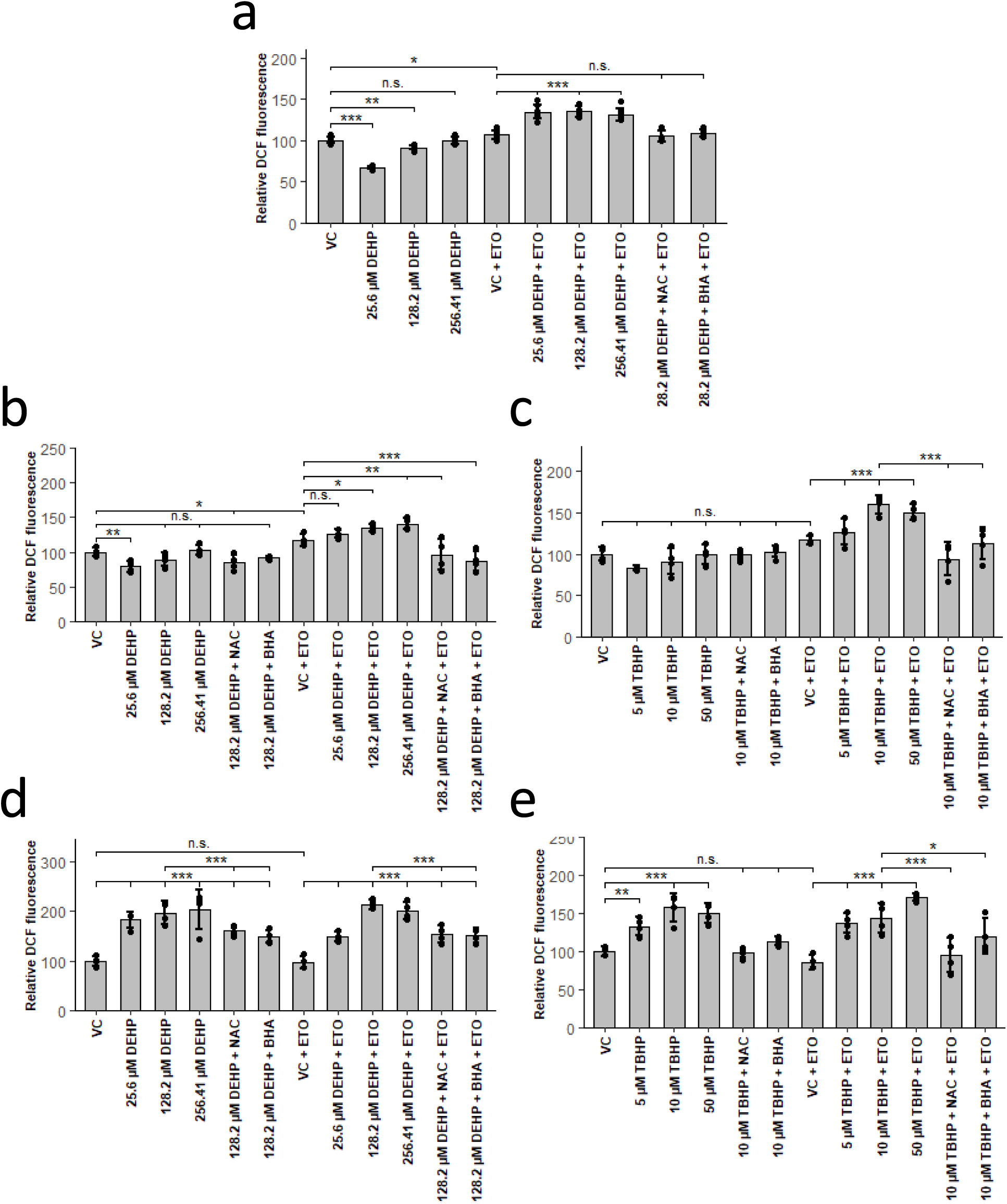
Enhanced ROS quenching capability depends on active fatty acid oxidation in neutrophils and other tissues. (a,b,d) Relative DCF fluorescence in neutrophils (a), HepG2/C3A (b) and HUVECs (d), with or without preincubation with 5 µM etomoxir (ETO) and 250 µM N-Acetylcysteine (NAC) or 10 µM butylated hydroxyanisole (BHA) for 2h, following 4h treatment with DEHP. (c,e) Relative DCF fluorescence in HepG2/C3A (c) and HUVECs (e), with or without preincubation with 5 µM etomoxir (ETO) and 250 µM N-Acetylcysteine (NAC) or 10 µM butylated hydroxyanisole (BHA) for 2h, following 4h treatment with tert-Butyl hydrogenperoxide (TBHP). Data are represented as mean ± SEM. Significance was assessed by one-way ANOVA (*p<0.05; **p<0.01; ***p<0.001, n.s. not significant).

As this mechanism should be generally present in cells owning active FAO, we tested if this hypothesis also holds true in other tissues. We therefore exposed HUVECs, generating little amounts of ATP via FAO, as well as HepG2/C3A cells, generating high amounts of ATP via FAO, to rising concentrations of DEHP in presence or absence of etomoxir (Dagher et al., 2001; Grünig et al., 2018). In line with our hypothesis, HepG2/C3A cells only showed increased ROS levels after pre-treatment with etomoxir (Fig. 5 b), whereas ROS increase in HUVECs was independent of etomoxir (Fig. 5 d). As in both cell lines stimulation of DCF fluorescence could be reduced by NAC and BHA, an underlying increase of ROS is evident. We further tested if our hypothesis also holds true for tert-butyl hydroperoxide (TBHP), another prominent ROS-inducer. Strikingly, usage of TBHP resulted in patterns similar to those observed with DEHP (Fig. 5 c,e). These results, therefore, clearly demonstrate that effects from DEHP are mediated via generation of ROS and are highly dependent on FAO activity. As this relation could also be observed in other cell lineages, we conclude this to be a general mechanism.

Taken together, our results show that DEHP treatment results in increased oxidative stress, in turn causing increased flux to NADPH-generating PPP. This, in turn, leads to ATP depletion in glycolytic active lineages, ultimately resulting in apoptosis. In lineages with active FAO, however, ATP generation is independent of glucose, resulting in significantly higher resistance against ROS-inducing compounds (see model, Fig. 6).

**Figure 6.**
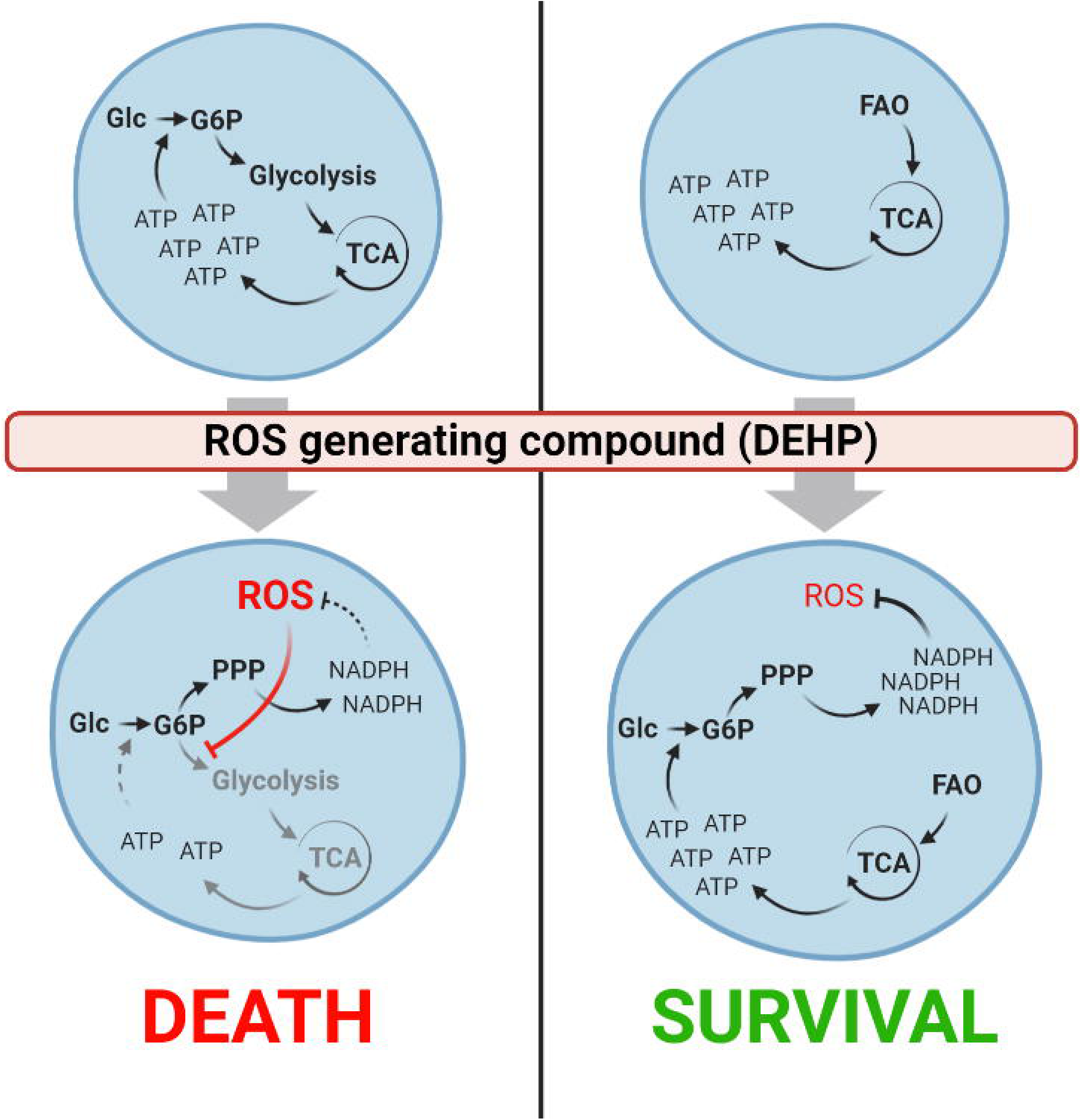
Active fatty acid oxidation enhances ROS quenching capability. Model depicting the critical role of active FAO in providing sufficient ATP levels used for glucose phosphorylation, resulting in active PPP. ROS induces a redirection of glycolytic flux through PPP, leading to lower levels of ATP in cells without active FAO (left). Lower levels of ATP, in turn, result in lower glucose phosphorylation and lower NADPH generation, ultimately leading to cell death. In cells with active FAO, ATP is generated independent of glycolysis, leading to higher levels of NADPH and thus ROS quenching capability (right).

## Discussion

Here we demonstrate, that DEHP selectively disturbs myeloid hematopoiesis in a concentration dependent manner. As shown, from the three lineages, erythroid and dendritic cell differentiation was considerably lowered by DEHP, neutrophil differentiation, however, was enhanced. Global metabolic profiling revealed a reduced glycolysis, glutaminolysis and polyamine synthesis in both lineages, as well as enhanced apoptosis. We here could show that the mode of action of DEHP is mediated by ROS generation, leading to fluxes shifting from glycolysis to PPP. This in turn leads to lowered levels of ATP in glycolytic active cell lines, which then again lowers PPP, ultimately resulting in apoptosis. In neutrophils, however, ATP is provided from FAO, preventing this fatal loop. We could further demonstrate that this relation also holds true in other tissues, as well as with other ROS-inducing compounds. Taken together, this relation seems to be a general mechanism, rendering tissues with active FAO more resistant against ROS inducing compounds.

To date, several different mechanisms, involved in ROS quenching, are known and can be classified as direct action of anti-oxidative enzymes, e.g. catalase, or production of reducing agents, e.g. glutathione (Birben et al., 2012; Lee and Paull, 2020; Snezhkina et al., 2020). Indeed, as NADPH acts as a major donor of reductive potential, NADPH-generating pathways (e.g. PPP) are most recognised to be involved in ROS-quenching. Here, redirection of glycolytic flux through PPP, as well as acceleration of glycolysis and PPP are known to be antioxidative mechanisms in response to elevated ROS levels (Snezhkina et al., 2020). An increase in FAO activity in healthy tissues, however, is mostly associated by an increase of ROS levels (Cheng et al., 2020; Kajihara et al., 2017; Lee and Paull, 2020; Xie et al., 2020). As a matter of fact, involvement of FAO as protective mechanism against ROS, is a quite well-known concept in cancer cells, resulting in new approaches for tumour therapy (Pike et al., 2011; Stoykova and Schlaepfer, 2019; Tabe et al., 2020). To our knowledge, the relation between distinctive responses to ROS-inducing xenobiotics and active FAO in healthy tissues surprisingly only receives little attention. As we could demonstrate, that FAO activity is the underlying cause for strongly differing effects of DEHP on hematopoietic lineages, we recommend that future toxicological studies should also address initial metabolic fluxes in their experimental systems. By this, differing responses to the same stimulus observed in different cell lineages, may be resolvable and lead to a better comparability of the different toxicological systems.

Considering high levels (up to 123.1 µg/ml) of DEHP exposure to infants treated in intensive care units, these results are actually highly relevant. As DEHP is known as immune adjuvant and exposure is linked to onset of asthma, we further provide a molecular mechanism of DEHP as a cause. Of note, ROS production was shown to be crucial for emergency granulopoiesis, which we believe to be the cause for enhanced neutrophil maturation, observed here (Kwak et al., 2015). In fact, increased counts of neutrophils have already been observed in bronchoalveolar lavage after DEHP administration in mice, providing further evidence *in vivo* (Larsen et al., 2007; You et al., 2014). In a nutshell, exposure to DEHP is likely to disturb hematopoiesis in humans; accordingly appropriate alternative plasticizers should replace DEHP as soon as possible. In addition, the herein observed results regarding DEHP should also, at least partially, be transferrable to other ROS-generating xenobiotics. Effects on hematopoiesis by such compounds should, therefore, be considered in future studies.

## Materials and Methods

### Reagents and materials

Human cord blood CD34^+^ cells were purchased from STEMCELL Technologies GmbH (Cologne, Germany). HepG2/C3A (ATCC CRL-10741) were purchased from American Type Culture Collection (Manassas, USA). StemPro-34 SFM Media, IMDM and valproic acid were from Fisher Scientific (Wien, Austria). Fms-related tyrosine kinase 3 ligand (Flt-3L), interleukin-3 (IL-3), interleukin-6 (IL-6), interleukin-15 (IL-15), thrombopoietin (TPO), stem cell factor (SCF) and erythropoietin (EPO) were purchased from GeneScript (Piscataway, NJ). BML-210 and stemregenin 1 (SR-1) were from ApexBio Technology LCC (Houston, TX). L-Glutamine (as Alanyl-L-Glutamine) was from Carl Roth GmbH+Co.KG (Karlsruhe, Germany). Di-2-ethylhexly phthalate (DEHP, analytical grade), Human Umbilical Vein Endothelial Cells (HUVEC), Endothelial Cell Growth Medium, DMEM (without phenol red) and human male AB serum were from Sigma-Aldrich Chemie GmbH (Munich, Germany). H_2_DCFDA, Rosiglitazone, fenofibrate, etomoxir, GW9662 and GW6471 were from VWR International GmbH (Bruchsal, Germany). Sphingolipids, including sphingosines, ceramides and sphingomyelines were purchased from Avanti Polar Lipids Inc. (Alabaster, Al, USA). All antibodies were purchased from Beckton Dickinson GmbH (Heidelberg, Germany).

Human CD34^+^ cell culture

Human cord blood CD34^+^ cells were expanded in Stempro-34 SFM media containing 2 mM L-glutamine, 100 ng/ml SCF, 100 ng/ml Flt-3L, 50 ng/ml IL-6, 40 ng/ml TPO, 1 µM SR-1, 0.1 µM BML-210 and 0.2 mM valproic acid (expansion media) as reported previously ^25^. Erythroid differentiation was performed in Stempro-34 SFM media containing 2 mM L-glutamine, 100 ng/ml SCF, 100 ng/ml Flt-3L, 50 ng/ml EPO and 20 ng/ml IL-3 as reported previously ^25^. Dendritic cell differentiation was done in Stempro-34 SFM media containing 2 mM L-glutamine, 10 ng/ml SCF, 10 ng/ml TPO and 10 ng/ml IL-3 as reported previously ^25^. Neutrophil differentiation was achieved in IMDM containing 10% fetal bovine serum, 50 ng/ml SCF and 50 ng/ml IL-15 as reported previously ^25^. Different concentrations of DEHP, solved in ethanol, pure ethanol (as vehicle control) or indicated agonists and antagonists were added as indicated at the beginning of each differentiation and held constant during the experiment, the added volume did not exceed 0.1% (v/v). Cell viability was routinely assessed using the trypan blue exclusion method.

### HUVEC and HepG2/C3A cell culture

Human Ubilical Vein Endothelial Cells were cultured in Endothelial Cell Growth Medium 3 at 37°C in a humidified atmosphere (5%CO_2_/95% air). HepG2/C3A cells were cultured in DMEM (without phenol red), supplemented with 10% fetal bovine serum, 100 U/ml penicillin and 100 mg/ml streptomycin 3 at 37°C in a humidified atmosphere (5%CO_2_/95% air). Cells were seeded into 96-well plates at a density of 4×10^4^ cells/well and left for attachment overnight prior the experiments.

### Flow cytometry

All flow cytometry data were acquired on a CyFlow Cube 8 flow cytometer (Sysmex) and analysed using FCS Express software (De Novo Software). Cells were washed twice with dPBS and blocked with 10% human male AB serum for 15 min at 4°C as reported previously ^58^. The cells were then double-stained with corresponding anti-human antibodies, using concentrations and incubation times according to the manufacturers instructions. After incubation, cells were washed twice dPBS and immediately analysed.

### Caspase 3/7 activation measurements

Erythrocytes were differentiated in presence of indicated concentrations of DEHP for 2 days, dendritic cells were differentiated for 3 days. Staurosporine was added for 18h. Prior the assay, cell solutions were diluted to 1 × 10^5^ cells/ml, using pre-warmed media. Caspase 3/7 activation was measured using the Apo-ONE Homogenous Caspase-3/7 Assay (Promega), following the manufacturers protocol. Cells were incubated with reagent for 1 h at room temperature.

### ATP quantification

Absolute ATP levels per 10^5^ cells were quantified using the ATP Detection Assay Kit – Luminescence (Cayman Chemical). In brief, lineages were differentiated in presence of DEHP for 2 days in case of erythrocytes and 3 days in case of dendritic cells and neutrophils. Cells then were washed twice with PBS and resuspended at 1×10^5^ cells per 167 µl ATP detection sample buffer. Cell lysis and ATP quantitation was performed following the manufacturers protocol.

### NADPH quantification

NADPH levels per 5×10^5^ cells were quantified using the NADPH Assay Kit (Colorimetric) (Abcam). In brief, lineages were differentiated in presence of DEHP for 2 days in case of erythrocytes and 3 days in case of dendritic cells and neutrophils. Cells then were washed twice with PBS at 4°C and resuspended at 1×10^6^ cells per 100 µl Lysis buffer. Cell lysis and NADPH quantitation was performed following the manufacturers protocol, lysates were incubated with reagent for 1 h at room temperature.

### H_2_DCFDC-Assay for detection of ROS

Relative cellular levels of reactive oxygen species (ROS) were determined by using 2’-7’-Dichlordihydrofluorescein-diacetate (H_2_ DCFDA) as described before ^6^. In brief, dendritic cells and neutrophils were differentiated for the indicated time spans in presence or absence of various concentrations of DEHP. Erythrocytes were differentiated for 2 days prior treatment with various concentrations of DEHP for 4 hours. HUVECs and HepG2/C3A were treated for 4 hours with DEHP or TBHP. ETO, NAC or BHA were added 2 h prior treatment with DEHP or TBHP. Following DEHP/TBHP treatment, H_2_DCFDA was added to each well, resulting in a final concentration of 20 µM and further incubated at 37°C for 30 min. Then cells were washed twice with PBS and resuspended at 1×10^5^ cells/100 µl. Measurement of oxidized dye, resulting in fluorescence, was carried out using a plate reader at Ex/Em: 485 nm/535 nm.

### Total RNA isolation and qPCR

Total RNA isolation was performed using the NucleoSpin RNA XS Kit (Macherey-Nagel GmbH &Co.KG). Gene specific primers were designed by use of Primer 3 software version 0.4.0 (MIT Center for Genome Research) (http://frodo.wi.mit.edu/cgi-bin/primer3/primer3_www.cgi) to obtain an annealing temperature of 56 °C and an amplicon length between 50 and 150 bp ^59^. Gene and species specificity were tested using NCBI nucleotide database, nucleotide blast, interrogation mode “blastn” (National Center for Biotechnology Information, Bethesda). First-strand complementary DNA (cDNA) synthesis was performed with 650 ng of isolated totalRNA (according to the manufacturer’s instructions). After adjusting total RNA volume to 11.5 µl, 1 µl (0.5 µg/µl) oligo-d(T)^18^-primer solution, 4 µl reaction-buffer 5x, 0.5 µl RiboLock® RNAse-inhibitor 40 U/mL, 1 µl Revert Aid Reverse Transcriptase® (RT), 2 µl 10 mM dNTP stock solution (Thermo Scientific) added, mixed and incubated 60 min at 42 °C. Enzyme was then inactivated by incubation at 70 °C for 10 min.

Real-time PCR was performed in a LightCycler 480 (Roche Diagnostics), using the LightCycler 480 SYBR Green I Master Kit (Roche Diagnostics) according to the manufacturers instructions. Raw data (ct-values) were extracted using LightCycler® 480 SW software® (version 1.5) (Roche Diagnostics). RPLP0 was chosen as reference gene for normalization of relative gene expression of each gene of interest. Relative gene expression was calculated using the Delta Delta CT method ^60^.

Each primer is characterized by the following characteristic features: symbol / gene / gene bank Accession / primer sequence forward (5’ to 3’) (fw), reverse (5’ to 3’) (rev) / amplicon size. Genes of interest displayed: HBB: hemoglobin subunit beta; NM_000518.4; fw: 5⍰-CTC GCT TTC TTG CTG TCC A-3’; rv: 5⍰-CAA GGC CCT TCA TAA TAT CCC C-3’, 88 bp. ELANE: elastase, neutrophil expressed; NM_001972.3; fw: 5⍰-CTG CGT GGC GAA TGT AAA CG-3’; rv: 5⍰-CGT TGA GCA AGT TTA CGG GG-3’, 140 bp. GPNMB: transmembrane glycoprotein NMB; NM_001005340.1 (transcript variant 1), NM_002510.2 (transcript variant 1); fw: 5⍰-CTG ATC TCC GTT GGC TGC TT-3’; rv: 5⍰-CTG ACC ACA TTC CCA GGA CT-3’; 110 bp. S100A8: S100 calcium binding protein A8; NM_002964.5; fw: 5’-GAT AAA GAT GGG CGT GGC AG-3’; 104 bp. Reference gene displayed: RPLP0: Ribosomal Protein, large, P0; NM_001002.3; fw: 5⍰-TGGCAATCCCTGACGCACCG-3⍰; rv: 5⍰-TGCCCATCAGCACCACAGCC-3⍰; 194 bp.

### Metabolite extraction

Cell pellets were washed twice with ice-cold PBS and the pellets were resuspended at 1×10^7^ cells/ml in -20°C cold ethanol containing 15% (v/v) 10 mM K_2_PO_4_, pH 7.5, sonicated for 3 min and frozen in liquid nitrogen for 30 seconds. The sonication-freeze cycle was repeated twice, afterwards, samples were centrifuged at 31,500 rcf at 2°C for 5 min and the supernatant was stored at -80°C until measurement.

### LC/MS-based metabolomics

The metabolome analyses were carried out with the AbsoluteIDQ p180 Kit (Biocrates Life Science AG) according to the manufacturers instructions. 20 µl cell lysates were mixed with isotopically labelled internal standards, derivatized with phenylisothiocyanate and extracted. LC-MS analysis, as well as FIA-ESI-MS/MS analysis was done using a 4000 QTRAP mass spectrometer (ABI Sciex) coupled to a NexeraXR HPLC (Shimadzu). A calibrator mix consisting of 7 different concentrations was used for calibration, also quality controls for 3 different concentration levels were included. The experiments were validated using the supplied MetIDQ software (Biocrates).

Further sphingolipids, including sphingosine, sphingosine-1-phosphate, C16-, C18-, C20-, C24-ceramide and corresponding dihydro-species, as well as lysosphingomyelin d18:1 and C16-sphingomyeline were quantified, using LC-MS. Briefly, 10 µl internal standard solution, consisting of 437.9 nM sphingosine d17:1, 684 nM sphingosine-1-phosphate d17:1, 477.1 nM C15 ceramide and 1.25 µM C24 Ceramide d17:1, was applied to 7 mm in diameter punches of 0.75 mm blotting paper, immobilized in an 96-well filter plate on top of a 96-deep well plate and dried under nitrogen flow (40 psi) for 20 minutes. Afterwards, 20 µl of calibrators or cell extracts were applied to each blotting paper and dried under nitrogen for 40 minutes.

300 µl of Methanol/methyl-tert-butyl-ether (MTBE) 80:20 was then applied to each well, sealed with a silicon mat and incubated for 30 min on a horizontal shaker at 450 rpm and 20 °C. The silicon mat then was removed, and extracts were transferred to the deep well plate by centrifugation at 500 rcf for 5 min at 4°C. The extracts were directly used for LC-MS measurement, the autosampler was set at a temperature of 10°C during the measurement.

Chromatographic separations on a NexeraXR HPLC (Shimadzu) were obtained under gradient conditions using a C8 Ultra-Inert HPLC column (50 mm L x 2.1 mm i.d., 3 µm particle size) (ACE), in combination with a C8 Ultra-Inert HPLC guard cartridge (10 mm L x 2.1 mm i.d., 3 µm particle size) (ACE), preheated at 50°C. The mobile phase consisted of eluent A (water with 0.4% formic acid) and eluent B (2-propanol with 0.4% formic acid). The gradient was as follows: From t=0 to 0.5 min A/B 80:20, followed from t=0.5 to 4.5 min by a linear gradient from 80:20 to 0:100, then from t=4.5 to 6 min 0:100 and finally from t=6 to 8 min A/B 80:20. The injection volume was 20 µl and the flow rate was set at 1 ml/min. A typical chromatogram is given in supplementary Fig. S2, the corresponding assay parameters for each analyte are given in supplementary Table 8. The 4000 QTRAP mass spectrometer (ABI Sciex) was operated in the positive ion mode with an electrospray voltage of 5000 V at 450°C, curtain gas at 25 psi, collision gas at 6 psi, nebulizing gas at 25 psi and auxiliary gas at 25 psi. All quadrupoles were working at unit resolution.

Typical precursor-to-product ion transitions (according to ^61^), with the corresponding parameters and retention times as stated in supplementary table 7 were used as quantifier for the scheduled multiple reaction monitoring (scheduled MRM), the retention time window was set at 60 seconds. Quantitation was performed with the MultiQuant V3.0.3 Software (ABI Sciex), using a 7-point calibration for each analyte, with the concentration ranges indicated in supplementary table 8. Ratios of analyte peak area and internal standard peak area were plotted against calibrator concentrations and calibration curves were calculated by least squares quadratic regression with 1/x weighting.

### Quantification and statistical analysis

#### Bioinformatic analysis

All data are represented as indicated in the corresponding legends. Statistical analysis of metabolomics data using an empirical Bayes approach was done by using the statistical software R ^62^ and the bioconductor package limma ^63^. Absolute log_2_ fold changes (logFC) above 0.5, with a corresponding FDR-adjusted p value below 0.05 where considered relevant. Furthermore, one-way ANOVA, where a p value smaller than 0.05 was considered as significant, was used for statistical analysis in indicated cases. Expansion rates are given as fold expansion, calculated as the resulting cell number divided by the initial cell number from the experiments. For metabolic pathway visualization, using Kyoto Encyclopedia of Genes and Genomes (KEGG)^64^ pathways as template, the software PathViso^65^ was used.

## Data availability

Significant metabolite datasets, generated in this study, can be found in the supplemental informations.

## Acknowledgments

This work was supported by a scholarship for Lars Kaiser from the Federal Ministry of Science, Research and Art of Baden-Württemberg; support by Steinbeis Center for Personalized Medicine (StZ1789) is gratefully acknowledged. M.J. thanks the German Research Foundation (DFG) under Germany’s Excellence Strategy (CIBSS – EXC-2189 – Project ID 390939984) for support. The authors also thank Markus Bläss for helpful discussions.

## Author Contributions

H.P.D., M.J. and R.C. were involved in study design. L.K. performed most experiments. I.Q. contributed to qPCR measurements. L.K. performed bioinformatic analysis. L.K., M.J. and H.P.D. wrote the paper, all authors commented on the manuscript.

## Supplemental Information titles and legends

**Figure S1**. Related to Figure 1-3; effects of DEHP are not mediated via modulation of PPARα/γ or fatty acid oxidation.

(a) Expansion rate of neutrophils treated with raising amounts of DEHP.

(b) Expansion of erythrocytes, treated with etomoxir, troglitazone, GW-9662, Fenofibrate, GW-6471 or DEHP.

(c) Expansion of erythrocytes in presence of DEHP alone, or DEHP in combination with different concentrations of etomoxir, GW-9662 or GW-6471.

(d) HBB/RPLP0 expression of erythrocytes, treated with etomoxir, troglitazone, GW-9662, Fenofibrate, GW-6471 or DEHP.

(e) HBB/RPLP0 expression of erythrocytes in presence of DEHP alone, or DEHP in combination with different concentrations of etomoxir or GW-9662. From samples, treated with DEHP and GW-6471, no total RNA could be extracted.

(f) Expansion of dendritic cells, treated with etomoxir, troglitazone, GW-9662, Fenofibrate, GW-6471 or DEHP.

(g) GPNMB/RPLP0 expression of dendritic cells, treated with etomoxir, troglitazone, GW-9662, Fenofibrate, GW-6471 or DEHP.

(h) GPNMB/RPLP0 expression of dendritic cells in presence of DEHP alone, or DEHP in combination with different concentrations of etomoxir or GW-9662. From samples, treated with DEHP and GW- 6471, no total RNA could be extracted.

(i) Expansion of dendritic cells in presence of DEHP alone, or DEHP in combination with different concentrations of etomoxir, GW-9662 or GW-6471.

**Figure S2**. Typical chromatogram obtained during separation of further sphingolipid analysis. Peaks correspond to the following analytes: **1** - lysoSM (d18:1), **2** -Sphingosine (d17:1), **3** - Sphingosine-1-Phosphate (d17:1), **4** - Sphingosine (d18:1), **5** - Sphingosine-1-Phosphate (d18:1), **6** - Sphinganine (d18:0), **7** - Sphinganine-1-Phosphate (d18:0), **8** - Sphingomyelin (d18:1/16:0), **9** - C15 Ceramide, **10** - C16 Ceramide, **11** - C16 Dihydroceramide, **12** - C18 Ceramide, **13** - C18 Dihydroceramide, **14** - C20 Ceramide, **15** - C20 Dihydroceramide, **16** - C24 Ceramide (d17:1), **17** - C24 Ceramide, **18** - C24 Dihydroceramide.

**Table S1**. Significant metabolite changes in erythroid differentiation after 2 days. Given is the log_2_fold change of FDR significant metabolites.

**Table S2**. Significant metabolite changes in erythroid differentiation after 4 days. Given is the log_2_fold change of FDR significant metabolites.

**Table S3**. Significant metabolite changes in erythroid differentiation after 6 days. Given is the log_2_fold change of FDR significant metabolites.

**Table S4**. Significant metabolite changes in dendritic cell differentiation after 3 days. Given is the log_2_fold change of FDR significant metabolites.

**Table S5**. Significant metabolite changes in dendritic cell differentiation after 8 days. Given is the log_2_fold change of FDR significant metabolites.

**Table S6**. Significant metabolite changes in dendritic cell differentiation after 11 days. Given is the log_2_fold change of FDR significant metabolites.

**Table S7**. Analyte parameters for the applied ceramide quantification method.

**Table S8**. Summary of concentration ranges, LOD, LLOQ, recovery rates, intra-assay CV and inter-assay CV of the applied ceramide quantification method.

