## Supplemental Fig. 1,2 and Table 1-8 for "Active fatty acid oxidation defines the cellular response towards reactive oxygen species"

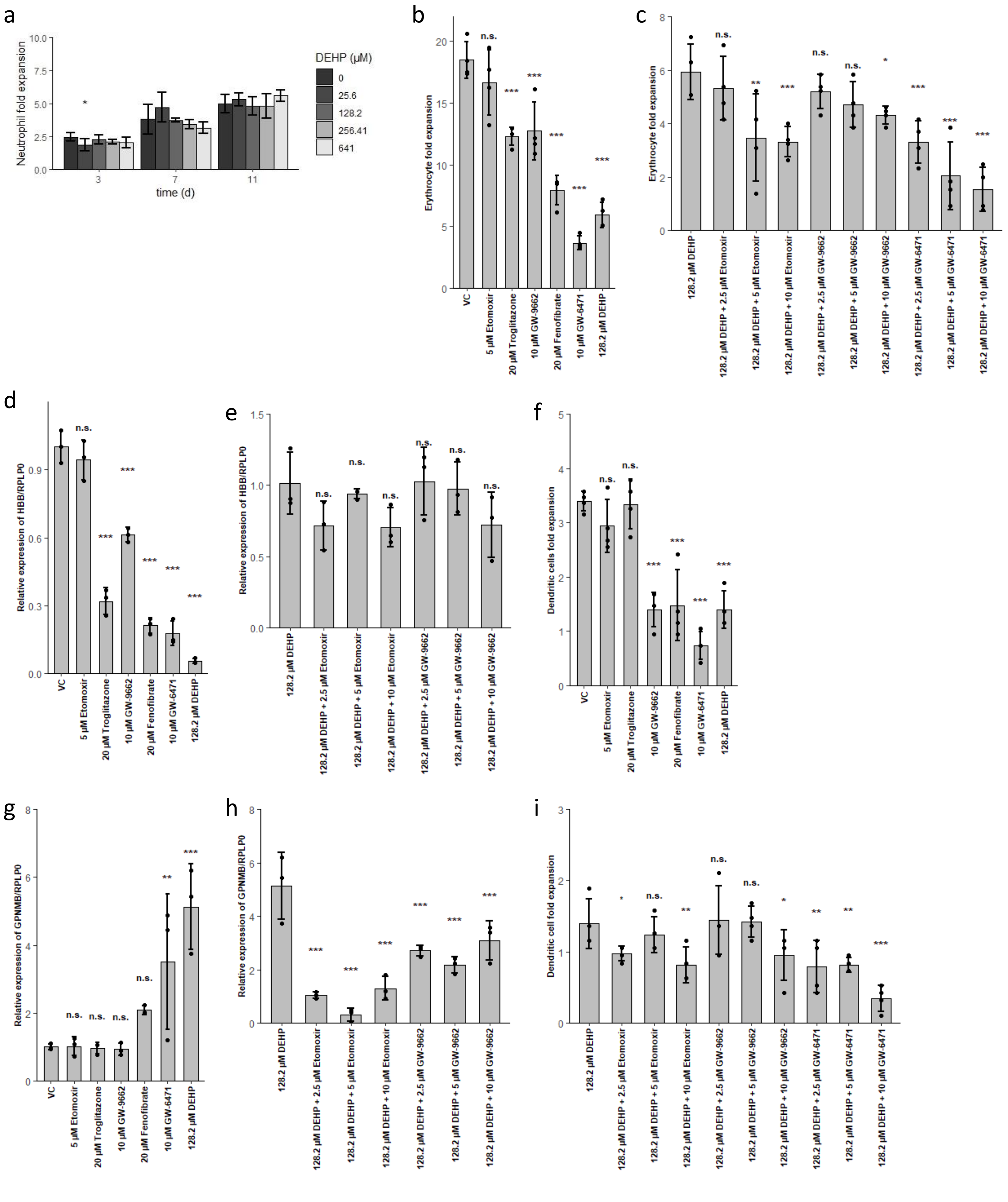


**Figure S1**. Related to Figure 1-3; effects of DEHP are not mediated via modulation of PPARα/γ or fatty acid oxidation.

(a) Expansion rate of neutrophils treated with raising amounts of DEHP.

(b) Expansion of erythrocytes, treated with etomoxir, troglitazone, GW-9662, Fenofibrate, GW-6471 or DEHP.
(c) Expansion of erythrocytes in presence of DEHP alone, or DEHP in combination with different concentrations of etomoxir, GW-9662 or GW-6471.
(d) HBB/RPLP0 expression of erythrocytes, treated with etomoxir, troglitazone, GW-9662, Fenofibrate, GW-6471 or DEHP.
(e) HBB/RPLP0 expression of erythrocytes in presence of DEHP alone, or DEHP in combination with different concentrations of etomoxir or GW-9662. From samples, treated with DEHP and GW-6471, no total RNA could be extracted.
(f) Expansion of dendritic cells, treated with etomoxir, troglitazone, GW-9662, Fenofibrate, GW-6471 or DEHP.
(g) GPNMB/RPLP0 expression of dendritic cells, treated with etomoxir, troglitazone, GW-9662, Fenofibrate, GW-6471 or DEHP.
(h) GPNMB/RPLP0 expression of dendritic cells in presence of DEHP alone, or DEHP in combination with different concentrations of etomoxir or GW-9662. From samples, treated with DEHP and GW-6471, no total RNA could be extracted.
(i) Expansion of dendritic cells in presence of DEHP alone, or DEHP in combination with different concentrations of etomoxir, GW-9662 or GW-6471.

*Table S1 Significant metabolite changes in erythroid differentiation after 2 days. Given is the $log_2$fold change of FDR significant metabolites.*

| Metabolite | 25.6 µM DEHP | 128.2 µM DEHP | 256.41 µM DEHP |
|---|---|---|---|
| *Glu* | 1.16 | | |
| *Gly* | 0.68 | 0.72 | |
| *His* | 0.55 | 0.63 | |
| *Ile* | 0.58 | 0.66 | |
| *Lys* | | 0.54 | |
| *Phe* | 0.47 | 0.52 | |
| *Pro* | 0.80 | 0.64 | 0.53 |
| *Thr* | 0.80 | | |
| *Val* | | 1.09 | |
| *Spermidine* | 0.80 | 0.58 | 0.48 |
| *Spermine* | 1.16 | 1.50 | 1.37 |
| *Taurine* | 2.13 | 1.84 | |
| *lysoPC.a.C18.1* | | 0.94 | |
| *lysoPC.a.C18.2* | | 1.87 | |
| *lysoPC.a.C20.4* | | | 1.27 |
| *lysoPC.a.C26.0* | | 0.69 | 0.92 |
| *lysoPC.a.C28.0* | 0.84 | | |
| *lysoPC.a.C28.1* | 0.59 | | |
| *PC.aa.C28.1* | 0.86 | | |
| *PC.aa.C30.0* | 1.46 | | 1.11 |
| *PC.aa.C30.2* | 1.00 | | |
| *PC.aa.C32.0* | 1.66 | 1.19 | 1.49 |
| *PC.aa.C32.1* | 1.23 | 0.88 | 0.89 |
| *PC.aa.C32.2* | 0.96 | 0.67 | 0.53 |
| *PC.aa.C32.3* | 0.81 | 0.47 | |
| *PC.aa.C34.1* | 1.29 | | |
| *PC.aa.C34.2* | 0.91 | 0.76 | |
| *PC.aa.C34.3* | 0.83 | 0.64 | 0.45 |
| *PC.aa.C34.4* | 0.75 | 0.46 | |
| *PC.aa.C36.1* | 1.19 | | |
| *PC.aa.C36.2* | 0.89 | | |
| *PC.aa.C36.3* | 0.83 | 0.66 | |
| *PC.aa.C36.4* | 0.96 | 0.77 | |
| *PC.aa.C36.5* | 0.77 | 0.47 | |
| *PC.aa.C36.6* | 0.86 | 0.65 | 0.44 |
| *PC.aa.C38.0* | 1.08 | 1.06 | 0.93 |
| *PC.aa.C38.3* | 0.87 | | |
| *PC.aa.C38.4* | 0.94 | | |
| *PC.aa.C38.5* | 0.70 | 0.39 | |
| *PC.aa.C38.6* | 0.80 | 0.78 | |
| *PC.aa.C40.2* | 1.09 | | |
| *PC.aa.C40.3* | 0.89 | | |
| *PC.aa.C40.4* | 0.94 | | |
| *PC.aa.C40.5* | 0.70 | 0.36 | |
| *PC.aa.C40.6* | 0.61 | 0.36 | |
| *PC.aa.C42.1* | 1.13 | 1.06 | 0.82 |
| *PC.aa.C42.2* | 0.62 | | |
| *PC.aa.C42.4* | 0.84 | | |

| | | | |
|---|---|---|---|
| *PC.aa.C42.5* | 1.12 | 0.85 | 0.76 |
| *PC.aa.C42.6* | 0.66 | 0.40 | |
| *PC.ae.C30.0* | 1.13 | 0.88 | 1.06 |
| *PC.ae.C30.1* | 1.10 | 0.94 | 0.93 |
| *PC.ae.C32.1* | 1.26 | 0.92 | 1.03 |
| *PC.ae.C32.2* | 1.16 | 0.86 | 1.00 |
| *PC.ae.C34.0* | 1.36 | 0.88 | 1.08 |
| *PC.ae.C34.1* | 1.33 | 0.98 | |
| *PC.ae.C34.2* | 1.12 | 0.98 | 0.98 |
| *PC.ae.C34.3* | 1.20 | 1.21 | 1.17 |
| *PC.ae.C36.1* | 1.22 | | |
| *PC.ae.C36.2* | 0.88 | 0.67 | |
| *PC.ae.C36.3* | 1.13 | 1.08 | 1.00 |
| *PC.ae.C36.4* | 1.30 | 1.07 | 1.07 |
| *PC.ae.C36.5* | 1.48 | 1.31 | 1.15 |
| *PC.ae.C38.0* | 0.70 | 0.38 | |
| *PC.ae.C38.1* | 0.94 | | |
| *PC.ae.C38.2* | 0.95 | | |
| *PC.ae.C38.3* | 0.98 | 0.78 | |
| *PC.ae.C38.4* | 1.22 | 1.02 | 0.98 |
| *PC.ae.C38.5* | 0.95 | 0.78 | 0.60 |
| *PC.ae.C38.6* | 1.34 | 1.25 | 1.13 |
| *PC.ae.C40.1* | 0.72 | 0.54 | |
| *PC.ae.C40.2* | 0.95 | | |
| *PC.ae.C40.3* | 0.86 | 0.75 | |
| *PC.ae.C40.4* | 0.95 | | |
| *PC.ae.C40.5* | 0.98 | 0.84 | 0.65 |
| *PC.ae.C40.6* | 1.05 | 0.95 | 0.77 |
| *PC.ae.C42.1* | 0.53 | | |
| *PC.ae.C42.2* | 0.94 | 0.64 | 0.50 |
| *PC.ae.C42.3* | 0.76 | 0.66 | 0.71 |
| *PC.ae.C44.3* | 0.70 | | |
| *PC.ae.C44.6* | | 0.70 | |
| *SM..OH..C16.1* | 0.99 | | |
| *SM..OH..C22.1* | | 1.20 | |
| *SM..OH..C24.1* | 1.04 | 0.97 | |
| *SM.C16.0* | 1.04 | | |
| *SM.C18.1* | 0.79 | | |
| *SM.C20.2* | 0.65 | 0.85 | |
| *SM.C22.3* | 0.80 | 0.85 | |
| *SM.C24.0* | 1.14 | | |
| *SM.C24.1* | 1.16 | | |
| *SM.C26.1* | 1.37 | 1.02 | |
| *H1* | 0.69 | 1.00 | |
| *S1P* | | | -3.30 |

*Table S2 Significant metabolite changes in erythroid differentiation after 4 days. Given is the $log_2$fold change of FDR significant metabolites.*

| Metabolite | 25.6 µM DEHP | 128.2 µM DEHP | 256.41 µM DEHP |
|---|---|---|---|
| *Asp* | | -1,69 | -1,93 |
| *Gln* | | -0,71 | |
| *Glu* | | -1,09 | |
| *Thr* | | -0,61 | |
| *Putrescine* | | -0,81 | -1,24 |
| *Spermidine* | -0,57 | -0,77 | -0,53 |
| *Taurine* | | -1,28 | |
| *C5.1* | | -0,78 | |
| *C14* | | -0,54 | |
| *lysoPC.a.C20.4* | | 0,97 | |
| *lysoPC.a.C28.0* | | -1,12 | |
| *PC.aa.C28.1* | | -1,22 | |
| *PC.aa.C30.2* | | -1,44 | |
| *PC.aa.C32.2* | | -0,55 | |
| *PC.aa.C34.1* | | -1,48 | |
| *PC.aa.C34.2* | | -0,61 | |
| *PC.aa.C34.3* | | -0,55 | |
| *PC.aa.C36.1* | | -1,89 | |
| *PC.aa.C36.2* | | -1,02 | -0,88 |
| *PC.aa.C36.3* | | -0,63 | |
| *PC.aa.C36.4* | | -0,95 | |
| *PC.aa.C38.3* | | -1,10 | |
| *PC.aa.C38.4* | | -1,00 | |
| *PC.aa.C38.5* | | -0,71 | -0,64 |
| *PC.aa.C40.3* | | -1,01 | |
| *PC.aa.C40.4* | | -1,12 | -0,97 |
| *PC.aa.C40.5* | | -0,60 | |
| *PC.aa.C42.4* | | -0,75 | |
| *PC.ae.C34.0* | | -1,33 | |
| *PC.ae.C36.1* | | -1,37 | |
| *PC.ae.C36.2* | | -0,58 | |
| *PC.ae.C36.5* | | 0,54 | |
| *PC.ae.C38.2* | | -0,89 | |
| *PC.ae.C38.3* | | -0,99 | |
| *PC.ae.C38.6* | | 0,51 | |
| *PC.ae.C40.2* | | -1,13 | |
| *PC.ae.C40.3* | | -0,90 | |
| *PC.ae.C40.4* | | -0,99 | -0,74 |
| *SM..OH..C16.1* | | -1,87 | -1,73 |
| *SM..OH..C22.1* | | -1,19 | |
| *SM..OH..C22.2* | | -1,72 | |
| *SM.C16.0* | | -1,75 | |
| *SM.C16.1* | | -0,89 | |
| *SM.C18.0* | | -1,99 | -1,44 |
| *SM.C18.1* | | -1,16 | -1,07 |
| *SM.C22.3* | | -0,96 | |
| *SM.C24.0* | | -1,23 | |
| *SM.C24.1* | | -1,64 | |

| | | | |
|---|---|---|---|
| *SM.C26.0* | | -1,00 | |
| *SM.C26.1* | | -2,24 | |
| *H1* | 0,74 | 0,30 | |
| *S1P* | -3.36 | -2.54 | -3.29 |
| *dhSphd18.0* | -1.63 | -1.65 | -1.57 |
| *C20Ceramide* | | | -1.26 |
| *C24Ceramide* | | -1.08 | |
| *C18DHCer* | | -1.00 | |

*Table S3 Significant metabolite changes in erythroid differentiation after 6 days. Given is the $log_2$fold change of FDR significant metabolites.*

| Metabolite | 25.6 µM DEHP | 128.2 µM DEHP | 256.41 µM DEHP |
|---|---|---|---|
| *Asp* | -1,16 | -1,45 | |
| *Putrescine* | -1,32 | -1,53 | |
| *Spermidine* | -0,61 | -0,57 | -0,43 |
| *Spermine* | | 0,36 | 0,96 |
| *C6..C4.1.DC.* | | -0,53 | |
| *C14.2* | | -0,98 | |
| *lysoPC.a.C16.1* | | | 0,86 |
| *lysoPC.a.C18.1* | | | 0,66 |
| *lysoPC.a.C18.2* | 1,37 | 1,53 | 1,53 |
| *lysoPC.a.C20.3* | | 0,83 | 0,90 |
| *lysoPC.a.C20.4* | 1,19 | 1,15 | 0,98 |
| *PC.aa.C30.2* | -0,74 | -0,57 | |
| *PC.aa.C32.0* | | | 1,21 |
| *PC.aa.C32.2* | -0,38 | -0,39 | -0,54 |
| *PC.aa.C32.3* | -0,31 | -0,45 | -0,92 |
| *PC.aa.C34.1* | -0,64 | | |
| *PC.aa.C34.3* | -0,53 | -0,63 | -0,77 |
| *PC.aa.C34.4* | | | -0,62 |
| *PC.aa.C36.1* | -1,06 | -0,92 | |
| *PC.aa.C36.4* | -0,38 | -0,46 | -0,59 |
| *PC.aa.C36.6* | | | -0,45 |
| *PC.aa.C38.3* | -0,60 | -0,64 | -0,62 |
| *PC.aa.C40.4* | -0,70 | -0,82 | -0,97 |
| *PC.aa.C42.4* | | | -0,60 |
| *PC.aa.C42.6* | 0,54 | 0,54 | |
| *PC.ae.C30.0* | | | 0,97 |
| *PC.ae.C30.1* | | 0,75 | 0,98 |
| *PC.ae.C30.2* | 0,71 | 0,67 | |
| *PC.ae.C32.1* | | | 1,03 |
| *PC.ae.C32.2* | | 0,62 | 0,81 |
| *PC.ae.C34.1* | | | 0,78 |
| *PC.ae.C34.2* | 0,54 | 0,68 | 1,19 |
| *PC.ae.C34.3* | 0,72 | 0,94 | 1,09 |
| *PC.ae.C36.1* | -0,64 | -0,58 | |
| *PC.ae.C36.3* | 0,61 | 0,81 | 0,84 |
| *PC.ae.C36.4* | 0,44 | 0,50 | 0,63 |
| *PC.ae.C36.5* | 0,98 | 1,18 | 0,89 |

| | | | |
|---|---|---|---|
| *PC.ae.C38.3* | | -0,57 | -0,47 |
| *PC.ae.C38.6* | 0,75 | 0,66 | 0,44 |
| *PC.ae.C40.1* | 0,69 | 0,83 | 0,63 |
| *PC.ae.C40.3* | -0,87 | -0,84 | -0,67 |
| *PC.ae.C40.4* | -0,49 | -0,62 | -0,67 |
| *PC.ae.C42.1* | 0,60 | 0,55 | 0,25 |
| *PC.ae.C42.2* | 0,96 | 1,08 | 0,86 |
| *SM..OH..C16.1* | -0,84 | -0,89 | |
| *SM..OH..C22.2* | -0,61 | | |
| *SM.C16.0* | | | |
| *SM.C18.0* | -1,21 | -1,23 | -0,75 |
| *SM.C18.1* | -0,78 | -0,94 | -0,95 |
| *SM.C20.2* | -0,65 | -0,95 | -0,89 |
| *SM.C22.3* | -0,61 | -0,63 | -0,85 |
| *SM.C24.0* | -0,67 | -0,68 | |
| *SM.C24.1* | -0,70 | -0,63 | |
| *H1* | 0,31 | -0,27 | |
| *C24Ceramide* | -1.08 | -1.14 | -1.96 |
| *dhSphd18.0* | | | -1.69 |
| *C20Ceramide* | | | -1.20 |
| *C18Ceramide* | | | -1.04 |
| *C16DHCer* | | 1.16 | |

*Table S4 Significant metabolite changes in dendritic cell differentiation after 3 days. Given is the $log_2$fold change of FDR significant metabolites.*

| Metabolite | 25.6 µM DEHP | 128.2 µM DEHP | 256.41 µM DEHP |
|---|---|---|---|
| *Lys* | | | 0,55 |
| *PC.aa.C32.0* | | | 1,68 |
| *PC.aa.C38.5* | | | -0,63 |
| *PC.ae.C34.0* | | | 1,47 |
| *H1* | | | 1,20 |
| *dhSphd18.0* | | -1.07 | -1.21 |
| *C18Ceramide* | | | -1.05 |

*Table S5 Significant metabolite changes in dendritic cell differentiation after 8 days. Given is the $log_2$fold change of FDR significant metabolites.*

| Metabolite | 25.6 µM DEHP | 128.2 µM DEHP | 256.41 µM DEHP |
|---|---|---|---|
| *Arg* | 1,60 | 1,84 | 1,37 |
| *Asp* | | -0,90 | -1,71 |
| *Glu* | -0,80 | -0,97 | -1,63 |
| *His* | | 0,86 | |
| *Ile* | 0,80 | 0,73 | 0,46 |
| *Leu* | 0,67 | 0,64 | |
| *Lys* | 0,95 | 1,02 | 0,72 |
| *Met* | 0,31 | 0,52 | 0,22 |
| *Phe* | 0,76 | 0,76 | 0,60 |
| *Ser* | 1,64 | | |
| *Thr* | | | |
| *Tyr* | 0,88 | 0,87 | 0,67 |
| *Val* | 1,06 | 0,90 | |
| *Histamine* | -1,92 | -2,17 | -2,97 |
| *Spermidine* | -0,36 | -0,53 | -0,86 |
| *Spermine* | | -0,58 | -0,79 |
| *Taurine* | -3,12 | -3,46 | -3,37 |
| *C14.2.OH* | | 0,52 | |
| *C16.OH* | 1,09 | 1,18 | 1,42 |
| *C16.1.OH* | 1,09 | 1,11 | 1,29 |
| *lysoPC.a.C16.0* | | | |
| *lysoPC.a.C16.1* | | | |
| *lysoPC.a.C17.0* | 1,59 | | 1,21 |
| *lysoPC.a.C18.0* | | | |
| *lysoPC.a.C18.1* | | | |
| *lysoPC.a.C18.2* | 1,06 | | |
| *lysoPC.a.C20.3* | | | -0,57 |
| *lysoPC.a.C20.4* | | -0,96 | -1,79 |
| *lysoPC.a.C24.0* | | | |
| *lysoPC.a.C26.1* | | | -1,10 |
| *lysoPC.a.C28.0* | | -0,82 | -1,84 |
| *lysoPC.a.C28.1* | | -0,91 | -1,71 |
| *PC.aa.C24.0* | | -0,61 | -0,86 |
| *PC.aa.C28.1* | -1,21 | -2,16 | -3,01 |
| *PC.aa.C30.0* | | -1,27 | -2,50 |
| *PC.aa.C30.2* | -1,19 | -2,44 | -3,89 |
| *PC.aa.C32.0* | 1,03 | | -1,24 |
| *PC.aa.C32.1* | | -1,48 | -2,97 |
| *PC.aa.C32.2* | | -2,13 | -3,22 |
| *PC.aa.C32.3* | -0,69 | -2,40 | -3,26 |
| *PC.aa.C34.1* | | -1,38 | -2,57 |
| *PC.aa.C34.2* | | -1,62 | -2,77 |
| *PC.aa.C34.3* | | -1,64 | -2,99 |
| *PC.aa.C34.4* | | -1,96 | -3,11 |
| *PC.aa.C36.0* | | -0,38 | -0,52 |
| *PC.aa.C36.1* | -0,93 | -1,75 | -2,89 |
| *PC.aa.C36.2* | -1,07 | -2,08 | -2,79 |
| *PC.aa.C36.3* | -0,56 | -1,73 | -2,75 |

| | | | |
|---|---|---|---|
| *PC.aa.C36.4* | -0,58 | -1,79 | -3,10 |
| *PC.aa.C36.5* | | -1,74 | -2,93 |
| *PC.aa.C36.6* | | -1,09 | -2,03 |
| *PC.aa.C38.0* | -0,71 | -1,69 | -2,00 |
| *PC.aa.C38.1* | | | -2,13 |
| *PC.aa.C38.3* | | -1,54 | -2,58 |
| *PC.aa.C38.4* | -0,67 | -1,75 | -2,88 |
| *PC.aa.C38.5* | -0,90 | -2,11 | -3,12 |
| *PC.aa.C38.6* | -0,37 | -1,44 | -2,54 |
| *PC.aa.C40.2* | | -0,98 | -1,99 |
| *PC.aa.C40.3* | | -1,09 | -2,47 |
| *PC.aa.C40.4* | | -1,48 | -2,29 |
| *PC.aa.C40.5* | -0,55 | -1,56 | -2,57 |
| *PC.aa.C40.6* | -0,37 | -1,11 | -1,50 |
| *PC.aa.C42.1* | | | |
| *PC.aa.C42.4* | | -0,78 | -1,54 |
| *PC.aa.C42.5* | | -0,59 | -1,73 |
| *PC.aa.C42.6* | | -0,61 | -0,84 |
| *PC.ae.C30.0* | | -0,95 | -1,57 |
| *PC.ae.C30.1* | | -1,63 | -2,56 |
| *PC.ae.C30.2* | | -1,32 | -2,25 |
| *PC.ae.C32.1* | | -1,45 | -2,79 |
| *PC.ae.C32.2* | | -1,46 | -2,55 |
| *PC.ae.C34.0* | | -0,88 | -2,00 |
| *PC.ae.C34.1* | | -1,27 | -2,39 |
| *PC.ae.C34.2* | | -1,05 | -2,15 |
| *PC.ae.C34.3* | | -0,79 | -2,17 |
| *PC.ae.C36.0* | | -0,68 | -0,90 |
| *PC.ae.C36.1* | -0,71 | -1,61 | -2,50 |
| *PC.ae.C36.2* | -0,45 | -1,42 | -2,02 |
| *PC.ae.C36.3* | | -1,08 | -2,00 |
| *PC.ae.C36.4* | -0,56 | -2,10 | -3,46 |
| *PC.ae.C36.5* | | -1,61 | -3,12 |
| *PC.ae.C38.0* | | -0,64 | -1,06 |
| *PC.ae.C38.1* | -0,95 | -1,63 | -2,31 |
| *PC.ae.C38.2* | -0,86 | -1,72 | -1,81 |
| *PC.ae.C38.3* | -0,56 | -1,60 | -2,74 |
| *PC.ae.C38.4* | | -1,85 | -2,94 |
| *PC.ae.C38.5* | -0,74 | -2,25 | -3,23 |
| *PC.ae.C38.6* | | -1,72 | -2,96 |
| *PC.ae.C40.1* | | -0,95 | -1,80 |
| *PC.ae.C40.2* | | -1,65 | -2,66 |
| *PC.ae.C40.3* | | -1,81 | -2,23 |
| *PC.ae.C40.4* | -0,62 | -1,44 | -1,63 |
| *PC.ae.C40.5* | | -1,68 | -2,49 |
| *PC.ae.C40.6* | | -1,91 | -2,69 |
| *PC.ae.C42.1* | | -0,98 | -1,49 |
| *PC.ae.C42.2* | | -0,97 | -2,05 |
| *PC.ae.C42.3* | | -1,13 | -1,79 |
| *PC.ae.C44.3* | | -0,61 | |
| *PC.ae.C44.5* | -0,48 | -0,85 | -1,03 |

| | | | |
|---|---|---|---|
| *PC.ae.C44.6* | -0,72 | -1,17 | -1,08 |
| *SM..OH..C14.1* | -0,74 | -1,57 | -2,47 |
| *SM..OH..C16.1* | -1,30 | -2,49 | -3,89 |
| *SM..OH..C22.1* | | | -0,50 |
| *SM..OH..C22.2* | | -1,42 | -2,63 |
| *SM..OH..C24.1* | | -1,39 | -2,13 |
| *SM.C16.0* | -0,98 | -2,08 | -3,64 |
| *SM.C16.1* | -0,65 | -1,69 | -2,76 |
| *SM.C18.0* | -1,39 | -2,45 | -3,79 |
| *SM.C18.1* | -1,08 | -2,26 | -3,52 |
| *SM.C20.2* | | -1,63 | -3,25 |
| *SM.C22.3* | | -2,06 | -3,21 |
| *SM.C24.0* | | -0,97 | -2,29 |
| *SM.C24.1* | | -1,46 | -2,91 |
| *SM.C26.1* | | -1,01 | -1,91 |
| *H1* | 1,84 | 1,92 | 1,71 |
| *dhSphd18.0* | | -2.60 | -5.79 |
| *C20Ceramide* | | -1.15 | -2.51 |
| *Sphd18.1* | | | -1.17 |
| *C18Ceramide* | | | -1.02 |

*Table S6 Significant metabolite changes in dendritic cell differentiation after 11 days. Given is the $log_2$fold change of FDR significant metabolites.*

| **Metabolite** | **25.6 µM DEHP** | **128.2 µM DEHP** | **256.41 µM DEHP** |
|---|---|---|---|
| *Ala* | | 0.6 | |
| *Arg* | 2,04 | 1,14 | 1,86 |
| *Gln* | | | |
| *Glu* | -1,36 | -1,12 | -1,04 |
| *Gly* | | -0,90 | |
| *His* | 0,74 | | 0,86 |
| *Ile* | 0,68 | | 1,15 |
| *Leu* | | | 0,93 |
| *Lys* | 1,15 | | 1,39 |
| *Phe* | 0,75 | | 0,97 |
| *Pro* | | | 0,40 |
| *Thr* | | | |
| *Tyr* | 0,72 | | 0,89 |
| *Histamine* | -2,84 | -1,85 | -4,44 |
| *Spermidine* | -0,55 | -0,36 | -0,55 |
| *Spermine* | -1,32 | -0,83 | -1,46 |
| *C2* | -0,75 | | -0,77 |
| *C3.1* | | | -1,26 |
| *lysoPC.a.C16.0* | 0,81 | | 1,10 |
| *lysoPC.a.C16.1* | | | -1,22 |
| *lysoPC.a.C18.1* | | | -1,15 |
| *lysoPC.a.C20.4* | -1,36 | | -2,29 |
| *lysoPC.a.C28.0* | | | -1,82 |
| *lysoPC.a.C28.1* | -1,36 | | -3,41 |
| *PC.aa.C24.0* | | | -0,79 |

| | | | |
|---|---|---|---|
| *PC.aa.C28.1* | -2,66 | -1,98 | -3,75 |
| *PC.aa.C30.0* | -1,24 | | -2,12 |
| *PC.aa.C30.2* | -1,94 | | -3,68 |
| *PC.aa.C32.1* | -2,17 | | -3,62 |
| *PC.aa.C32.2* | -3,82 | -1,63 | -5,52 |
| *PC.aa.C32.3* | -4,12 | -2,17 | -5,35 |
| *PC.aa.C34.1* | | | -1,59 |
| *PC.aa.C34.2* | -2,39 | -1,36 | -3,67 |
| *PC.aa.C34.3* | -3,18 | -1,39 | -4,88 |
| *PC.aa.C34.4* | -3,54 | | -5,77 |
| *PC.aa.C36.1* | | | -1,73 |
| *PC.aa.C36.2* | -2,48 | -1,81 | -3,48 |
| *PC.aa.C36.3* | -2,46 | -1,38 | -3,95 |
| *PC.aa.C36.4* | -2,12 | | -3,84 |
| *PC.aa.C36.5* | -2,76 | | -4,97 |
| *PC.aa.C36.6* | -2,22 | | -3,57 |
| *PC.aa.C38.0* | -2,00 | -1,41 | -2,63 |
| *PC.aa.C38.1* | -1,56 | | -0,86 |
| *PC.aa.C38.3* | -1,84 | -1,24 | -2,96 |
| *PC.aa.C38.4* | -1,94 | | -3,52 |
| *PC.aa.C38.5* | -2,60 | | -4,54 |
| *PC.aa.C38.6* | -1,92 | | -3,16 |
| *PC.aa.C40.2* | -0,94 | | -1,31 |
| *PC.aa.C40.3* | -1,42 | -0,86 | -2,42 |
| *PC.aa.C40.4* | -2,12 | -1,07 | -2,98 |
| *PC.aa.C40.5* | -2,53 | | -4,50 |
| *PC.aa.C40.6* | -1,23 | -0,59 | -1,64 |
| *PC.aa.C42.1* | -1,02 | | |
| *PC.aa.C42.4* | -0,91 | | -1,58 |
| *PC.aa.C42.5* | | | -3,09 |
| *PC.aa.C42.6* | -0,76 | | -1,05 |
| *PC.ae.C30.0* | | | -1,48 |
| *PC.ae.C30.1* | -2,25 | | -4,42 |
| *PC.ae.C30.2* | -2,32 | -0,77 | -2,53 |
| *PC.ae.C32.1* | -1,65 | -1,35 | -3,11 |
| *PC.ae.C32.2* | -2,41 | -1,51 | -3,41 |
| *PC.ae.C34.1* | | | -1,92 |
| *PC.ae.C34.2* | -1,75 | -1,21 | -3,43 |
| *PC.ae.C34.3* | -1,89 | | -4,09 |
| *PC.ae.C36.1* | | | -1,87 |
| *PC.ae.C36.2* | -1,81 | -1,48 | -3,11 |
| *PC.ae.C36.3* | -1,72 | -1,07 | -3,80 |
| *PC.ae.C36.4* | -2,95 | -1,38 | -4,79 |
| *PC.ae.C36.5* | -2,45 | -0,92 | -4,63 |
| *PC.ae.C38.1* | -1,91 | -1,94 | -2,05 |
| *PC.ae.C38.2* | -1,41 | -1,49 | -2,08 |
| *PC.ae.C38.3* | -2,26 | -1,67 | -3,59 |
| *PC.ae.C38.4* | -2,51 | -1,36 | -3,78 |
| *PC.ae.C38.5* | -3,05 | -1,61 | -4,63 |
| *PC.ae.C38.6* | -2,64 | | -4,54 |
| *PC.ae.C40.1* | -0,84 | | -1,63 |

| | | | |
|---|---|---|---|
| *PC.ae.C40.2* | -1,50 | | -2,40 |
| *PC.ae.C40.3* | -2,15 | -2,07 | -3,15 |
| *PC.ae.C40.4* | -2,14 | -1,63 | -2,71 |
| *PC.ae.C40.5* | -2,71 | -1,40 | -4,75 |
| *PC.ae.C40.6* | -2,90 | -1,34 | -4,13 |
| *PC.ae.C42.1* | -0,97 | | -1,17 |
| *PC.ae.C42.2* | -1,44 | | -1,52 |
| *PC.ae.C42.3* | -1,60 | -1,64 | -2,36 |
| *PC.ae.C44.3* | | | -0,66 |
| *PC.ae.C44.5* | -1,07 | -0,74 | -1,06 |
| *PC.ae.C44.6* | -2,03 | -1,49 | -2,13 |
| *SM..OH..C14.1* | -2,24 | | -2,79 |
| *SM..OH..C16.1* | -1,81 | | -4,06 |
| *SM.C16.0* | -1,91 | -1,54 | -3,47 |
| *SM.C16.1* | -2,21 | -1,33 | -3,23 |
| *SM.C18.0* | -1,56 | -1,94 | -2,84 |
| *SM.C18.1* | -2,37 | -1,89 | -3,56 |
| *SM.C20.2* | | | -3,72 |
| *SM.C22.3* | | | -2,62 |
| *SM.C24.0* | | | -1,37 |
| *SM.C24.1* | | | -1,88 |
| *SM.C26.0* | | | -0,92 |
| *H1* | 2,21 | 0,76 | 2,51 |
| *Sphd18.1* | | -2.41 | -1.48 |
| *C20Ceramide* | | | -1.37 |

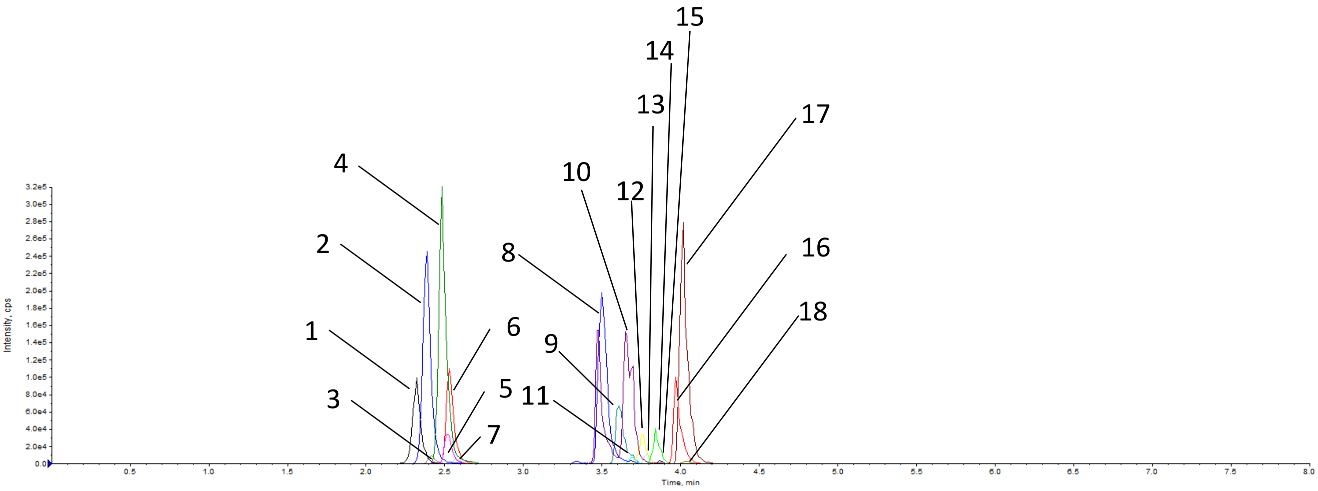


**Figure S2.** Typical chromatogram obtained during separation of further sphingolipid analysis.
Peaks correspond to the following analytes: **1** - lysoSM (d18:1), **2** -Sphingosine (d17:1), **3** - Sphingosine-1-Phosphate (d17:1), **4** - Sphingosine (d18:1), **5** - Sphingosine-1-Phosphate (d18:1), **6** - Sphinganine (d18:0), **7** - Sphinganine-1-Phosphate (d18:0), **8** - Sphingomyelin (d18:1/16:0), **9** - C15 Ceramide, **10** - C16 Ceramide, **11** - C16 Dihydroceramide, **12** - C18 Ceramide, **13** - C18 Dihydroceramide, **14** - C20 Ceramide, **15** - C20 Dihydroceramide, **16** - C24 Ceramide (d17:1), **17** - C24 Ceramide, **18** - C24 Dihydroceramide.

*Table S7 Analyte parameters for the applied ceramide quantification method.*

| Analyte | Retention time | m/z pair | Internal Standard | DP | CE | CXP |
|---|---|---|---|---|---|---|
| *Sphingosine* | 2.5 | 300.5/282.3 | Sphingosine(d17:1) | 41 | 19 | 6 |
| *Sphinganine* | 2.54 | 302.6/284.5 | Sphingosine(d17:1) | 76 | 23 | 24 |
| *Sphingosine-1-Phosphate* | 2.53 | 380.3/264.4 | Sphingosine-1-Phosphate(d17:1) | 71 | 21 | 18 |
| *Sphinganine-1-Phosphate* | 2.59 | 382.3/284.5 | Sphingosine-1-Phosphate(d17:1) | 50 | 22 | 25 |
| *C16 Ceramide* | 3.67 | 538.4/264.1 | C15 Ceramide | 70 | 30 | 15 |
| *C16 Dihydroceramide* | 3.70 | 540.9/284.5 | C15 Ceramide | 126 | 47 | 12 |
| *C18 Ceramide* | 3.76 | 567/264.4 | C15 Ceramide | 40 | 40 | 15 |
| *C18 Dihydroceramide* | 3.79 | 569/284.5 | C15 Ceramide | 70 | 45 | 14 |
| *C20 Ceramide* | 3.85 | 595.3/264.2 | C24 Ceramide (d17:1) | 70 | 45 | 15 |
| *C20 Dihydroceramide* | 3.88 | 597.3/284.5 | C24 Ceramide (d17:1) | 121 | 47 | 16 |
| *C24 Ceramide* | 4.02 | 650.8/264.2 | C24 Ceramide (d17:1) | 86 | 43 | 14 |
| *C24 Dihydroceramide* | 4.04 | 653.1/284.6 | C24 Ceramide (d17:1) | 126 | 45 | 16 |
| *Sphingomyelin (d18:1/16:0)* | 3.52 | 704/184.2 | None | 150 | 12 | 15 |
| *LysoSM (d18:1)* | 2.33 | 465.5/184.2 | None | 56 | 41 | 20 |

*Table S8 Summary of concentration ranges, LOD, LLOQ, recovery rates, intra-assay CV and inter-assay CV of the applied ceramide quantification method.*

| Analyte | Concentration range (nM) | $R^2$ (extracted calibrators) | LOD (S/N > 3) | LLOQ (S/N >10) | Recovery (standard) | Recovery (cell extract) | Intra-assay CV | Inter-Assay CV |
|---|---|---|---|---|---|---|---|---|
| *Sphingosine* | 1.67 - 835 | 0.97 | 3.34 | 3.34 | 78,57% | 84,07% | 18.15% | 15.31% |
| *Sphinganine* | 1.66 - 829 | 0.98 | 3.32 | 3.32 | 79,26% | 88,09% | 13.14% | 14.04% |
| *Sphingosine-1-Phosphate* | 2.64 – 1318 | 0.97 | 2.64 | 2.64 | 101,73% | 73,39% | 14.83% | 6.09% |
| *Sphinganine-1-Phosphate* | 13.1 – 1311 | 0.99 | 13.1 | 13.1 | 83,43% | 63,16% | 12.62% | 4.86% |
| *C16 Ceramide* | 1.86 – 930 | 0.98 | 1.86 | 1.86 | 96,26% | 76,12% | 12.00% | 22.57% |
| *C16 Dihydroceramide* | 1.85 – 926 | 0.96 | 1.85 | 1.85 | 103,16% | 19,09% | 17.35% | 14.71% |
| *C18 Ceramide* | 1.77 – 883 | 0.98 | 1.77 | 1.77 | 134,63% | 88,7% | 16.41% | 17.03% |
| *C18 Dihydroceramide* | 1.76 – 880 | 0.97 | 1.76 | 1.76 | 119,98% | 110,38% | 14.76% | 6.57% |
| *C20 Ceramide* | 1.68 – 842 | 0.94 | 1.68 | 1.68 | 109,62% | 83,98% | 20.02% | 30.56% |
| *C20 Dihydroceramide* | 1.68 – 839 | 0.93 | 1.68 | 10 | 97,17% | 107,41% | 22.31% | 10.85% |
| *C24 Ceramide* | 6.15 – 3076 | 0.95 | 6.15 | 6.15 | 71,17% | 61,19% | 16.86% | 6.98% |
| *C24 Dihydroceramide* | 1.53 – 767 | 0.98 | 1.53 | 30.76 | 66,1% | 51,95% | 30.45% | 37.67% |
| *Sphingomyelin (d18:1/16:0)* | 284.48 – 142242 | 0.97 | 284.48 | 284.48 | 84,14% | 135,42% | 13.01% | 15.45% |
| *LysoSM (d18:1)* | 2.15 - 1076 | 0.99 | 2.15 | 2.15 | 86,99% | 91,02% | 13.58% | 27.56% |
